# Single-cell transcriptomics of the naked mole-rat reveals unexpected features of mammalian immunity

**DOI:** 10.1101/597195

**Authors:** HG Hilton, ND Rubinstein, P Janki, AT Ireland, N Bernstein, KM Wright, D Finkle, B Martin-McNulty, M Roy, M Smith, DM Imai, V Jojic, R Buffenstein

## Abstract

Using single-cell transcriptional profiling we mapped the immune system of the naked mole-rat (*Heterocephalus glaber*), a small but long-lived and cancer-resistant subterranean rodent. Both splenic and circulating immune cells were examined in healthy young animals and following an infection-mimicking lipopolysaccharide challenge. Our study revealed that the naked mole-rat immune system is characterized by a high myeloid to lymphoid cell ratio that includes a novel, lipopolysaccharide responsive, granulocyte cell subset not found in the mouse. Conversely, we find that naked mole-rats do not have a cell subset that corresponds to natural killer cells as defined in other well-characterized mammalian species. Supporting this finding, we show that the naked mole-rat genome has not expanded any of the gene families encoding diverse natural killer cell receptors, which are the genomic hallmarks of species in which natural killer cells have been described. These unusual features suggest an atypical mode of immunosurveillance and a greater reliance on myeloid-biased innate immunity.

## Introduction

Unlike most mammals, the subterranean dwelling naked mole-rat (Rodentia; Ctenohystrica, *Heterocephalus glaber*) is cancer resistant, extremely long-lived relative to its body size and does not experience an exponential increase in risk of dying with increasing age (1,2). Moreover, they exhibit few, if any, signs of physiologic decline or age-associated molecular change (3–5). This suggests that they retain one or more biological features that allows them to maintain cellular and systemic homeostasis well beyond their age of sexual maturity and into their third decade of life (6). An unexplored possibility is that naked mole-rats have evolved features of systemic immunity that allow for enhanced cancer immunosurveillance or which prevent the age-associated increase in morbidity and mortality that is associated with weakened or altered immunity in other species (7,8).

The immune system comprises a network of specialized cells, tissues and organs that function in a highly coordinated way to mediate host defense against pathogens and malignantly transformed cells, as well as playing a role in the development, metabolism, and repair of tissues. Our understanding of these diverse roles has been informed through advances in microscopy and antibody-based flow cytometry that have allowed for the determination of the phenotype and function of the cells comprising the immune system at ever increasing resolution. However, an in-depth examination of the extent to which distinctive ecological niches have shaped the evolution of immune systems in non-model organisms has been restricted due to a lack of species-specific reagents with which immune cells can be isolated and characterized. Recent advances in single-cell RNA sequencing (scRNA-seq) have circumvented these traditional restrictions, allowing for the unbiased molecular characterization of the immune repertoire, including rare and novel cell subsets, in previously uncharacterized species (9).

In this study we investigated the immune system of the naked mole-rat using comparative single-cell transcriptomics and comparative genomics. We show that the spleen and circulating immune cell composition of the naked mole-rat exhibits major differences to that of the short-lived and cancer-prone mouse, with an inverted myeloid to lymphoid ratio, a novel lipopolysaccharide responsive granulocyte cell subset and no detectable natural killer (NK) cells. In addition, we found that the naked mole-rat does not have an expanded family of NK cell receptors that recognize MHC class I, suggesting that classical NK cell-mediated defense is not an important feature of naked mole-rat immunity or cancer resistance.

## Results

### Single-cell RNA-sequencing profiling of spleen from naked mole-rats and mice

Spleens were harvested from four healthy C57BL/6 mice (2-3 months old) and four healthy naked mole-rats (2 years old) and single cell suspensions were prepared from each organ (Figure 1a). We determined the gene expression in each spleen at single-cell resolution using droplet-based single-cell RNA sequencing (10X Genomics) (10). Each sample was run in duplicate, capturing between 5,495 and 15,930 cells from each mouse spleen and between 6,215 and 8,740 cells from each naked mole-rat spleen (Figure 1b and Tables S1-2). We analyzed a total of 39,088 and 29,949 cells from the mouse and naked mole-rat spleens respectively. The median number of genes detected per cell was 1,134 (range 177-5,635) and 737 (range 208-5,345) in the mouse and naked mole-rat respectively (Figure 1c and Tables S1-2), a disparity that likely arises due to differences in the genome annotations of each species. The median number of unique molecular identifiers (UMIs) detected per cell was 2,871 (range 242-42,756) and 1,854 (range 232-45,499) in the mouse and naked mole-rat respectively (Figure 1d and Tables S1-2).

**Figure 1:**
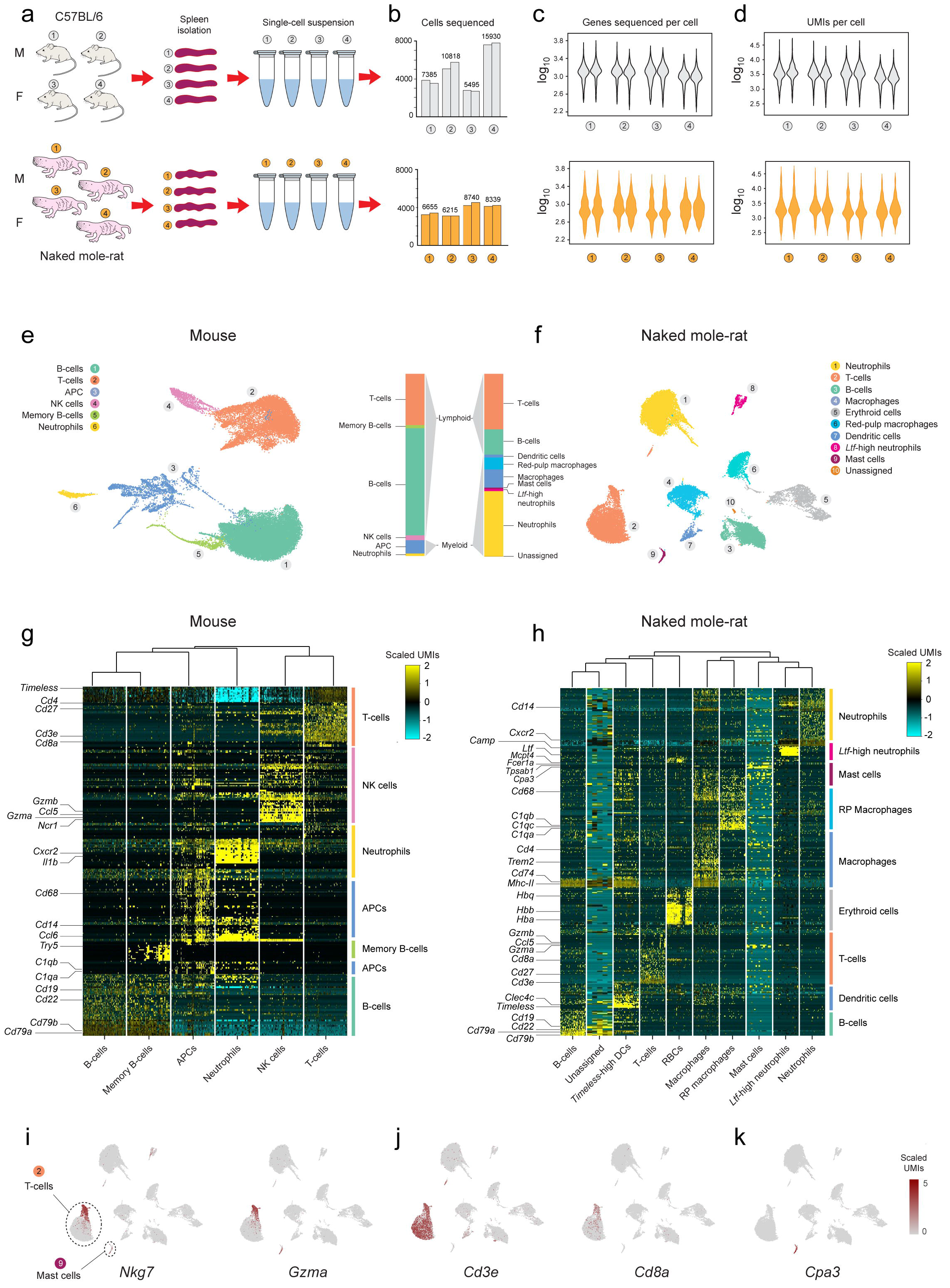
Single-cell transcriptional profiling of mouse and naked mole-rat spleens reveals major differences in immune cell populations. **a** - Schematic showing the workflow in which single-cell suspensions are derived from the spleens of four (two males [M], two females [F]) C57BL/6 mice (upper panel, grey) or four naked mole-rats (two males [M], two females [F]) (lower panel, orange). **b** - Bar chart showing the number of cells sequenced in duplicate from each of four mice (upper panel, grey) and four naked mole-rats (lower panel, orange). **c** - Violin plots showing the number of genes sequenced per cell from each of four mice (upper panel, grey) and four naked mole-rats (lower panel, orange). **d** - Violin plots showing the number of unique molecular identifiers (UMIs) sequenced per cell from each of four mice (upper panel, grey) and four naked mole-rats (lower panel, orange). **e** - Mouse spleen cell clusters visualized by uniform manifold approximation and projection (UMAP). Colors indicate clusters that correspond to the listed cell types. The relative proportions of each cell type are shown in a stacked bar chart to the right. **f** - Naked mole-rat spleen cell clusters visualized by UMAP. Colors indicate clusters that correspond to the listed cell types. The relative proportions of each cell type are shown in a stacked bar chart to the left. **g** - Heatmap showing the gene expression in mouse spleen cells in each of six clusters that correspond to the listed cell types. Selected marker genes are listed to the left. **h** - Heatmap showing the gene expression in naked mole-rat spleen cells in each of ten clusters that correspond to the listed cell types. Selected marker genes are listed to the left. Naked mole-rat cell clusters visualized by UMAP showing the relative expression of *Nkg7* and *Gzma* (**i**), *Cd3e* and *Cd8a* (**j**) and *Cpa3* (**k**).

### The mouse spleen is dominated by lymphoid lineage cells while the naked mole-rat spleen is dominated by myeloid lineage cells

We performed an iterative clustering approach (see *Methods*), in order to assign the cells into transcriptionally-defined clusters and compare the cellular heterogeneity of mouse and naked mole-rat spleens. In the mouse dataset, six clusters were identified in the first iteration of our clustering approach (Figures 1e, S1a and Table S1) (breaking down to a total of 26 clusters at convergence, Figures S1b, S2a and Table S1), whereas in the naked mole-rat dataset, ten clusters were identified in the first iteration (Figures 1f, S1c and Table S2) (breaking down to a total of 22 at convergence, Figures S1d, S2b and Table S2). In both species, clusters were demarcated by unique sets of highly and lowly expressed genes (Figures 1g-h and S2c-d respectively). Using differential-expression analyses (see *Methods*), we obtained a set of cluster-specific marker genes (Tables S3-4), which allowed us to annotate the cell types these clusters represent and hence reveal the major immune-cell subsets present in each species.

Consistent with previous reports (11,12), the mouse spleen is dominated by lymphoid lineage cells (~93% of the sequenced cells) comprising ~60% B-cells (marked by high expression of *Cd19* and *Cd79a* in clusters 1 and 5), ~30% T-cells (marked by high expression of *Cd3e* in cluster 2) and ~3% NK cells (marked by high expression of *Ncr1* in cluster 4). Myeloid cells form a small minority of the mouse splenic population, accounting for ~10% of the sequenced cells and comprising 8% antigen presenting cells (APCs) (marked by high expression of *Cd68*, *Ccl6* and *C1qa* in cluster 3) and ~2% neutrophils (marked by high expression of *Cxcr2* in cluster 6).

By contrast, lymphoid lineage cells comprise just 40% of the splenic population in naked mole-rats (25% T-cells with high expression of *Cd3e* in cluster 2 and 15% B-cells with high expression of *Cd79a* in cluster 3), while myeloid lineage cells account for ~60% of the sequenced cells. These cells comprise ~35% neutrophils (marked by high expression of *Cxcr2* in cluster 1) and ~15% macrophages (marked by high expression of *Cd68*, *C1qa* and *C1qb* in clusters 4 and 6). The remaining 10% of myeloid lineage cells comprise five rarer cell subsets, each accounting for 1-2% of the total cell repertoire. We have tentatively identified cluster 6 as red-pulp macrophages on the basis of their high expression of macrophage specific markers and their expression of *Vcam-1* (13). Also included in this myeloid cell subset we identified ~2% dendritic cells (high expression of *Clec4c* in cluster 7). Further distinguishing this dendritic-cell population is high expression of *Timeless* (Figure 1h), a gene more commonly associated with the regulation of circadian rhythm in drosophila (14). The function of *Timeless* in mammalian biology is less well understood but is thought to include the protection of telomeres from DNA damage (15). Mast cells (marked by high expression of *Cpa3*, *Mcpt4, Tpsab1 and Fcer1a* in cluster 9) comprise ~1% of the naked mole-rat spleen cell population. The naked mole-rat also contains a novel myeloid cell subset (cluster 8, proportion ~1% of sequenced cells) with a similar expression profile to neutrophils and mast cells (Figure 1h). This cell subset, which was identified in all four animals profiled, is defined by several marker genes including *lactotransferrin* (*Ltf*), *olfactomedin-4* (*Olfm4*), *cysteine-rich secretory protein-3* (*Crisp3*) and *cathelicidin* (*Camp*) (Figures 1h, S2b, S2d and Table S4). For simplicity we have termed these cells *Ltf-high neutrophils*. Thus, the naked mole-rat has an inverted myeloid to lymphoid ratio as compared to the mouse, driven predominantly by a reduction in the proportion of B-cells.

To confirm the inverted myeloid to lymphoid ratio observed between the mouse and naked mole-rat in our scRNA-seq data, which is clearly illustrated by the expression of *Cd14* in myeloid cells and *Cd19* and *Cd3e* in B- and T-lymphocytes respectively (Figure 2a), we performed a comparative histomorphological and in-situ hybridization (ISH) analysis in four spleens from each species. Formalin fixed paraffin-embedded tissue sections were stained with hematoxylin and eosin (H&E) or incubated with species-specific probes targeting *Cd14*, *Cd19* and *Cd3e*. The H&E stained mouse and naked mole-rat spleens showed major differences in splenic microanatomy, with the naked mole-rat having a comparatively reduced white pulp compartment, a larger red pulp compartment with a greater number of fibromuscular trabeculae that connect to the capsule and provide structural support and contractility to the spleen (Figure 2b). Within the white pulp, the marginal zone and follicles of the naked mole-rat (which comprise B-cells) are readily identifiable. By contrast, the periarteriolar lymphoid sheath (PALS) (which comprises the T-cell rich compartment in other species) is less prominent in the naked mole-rat than in the mouse.

**Figure 2:**
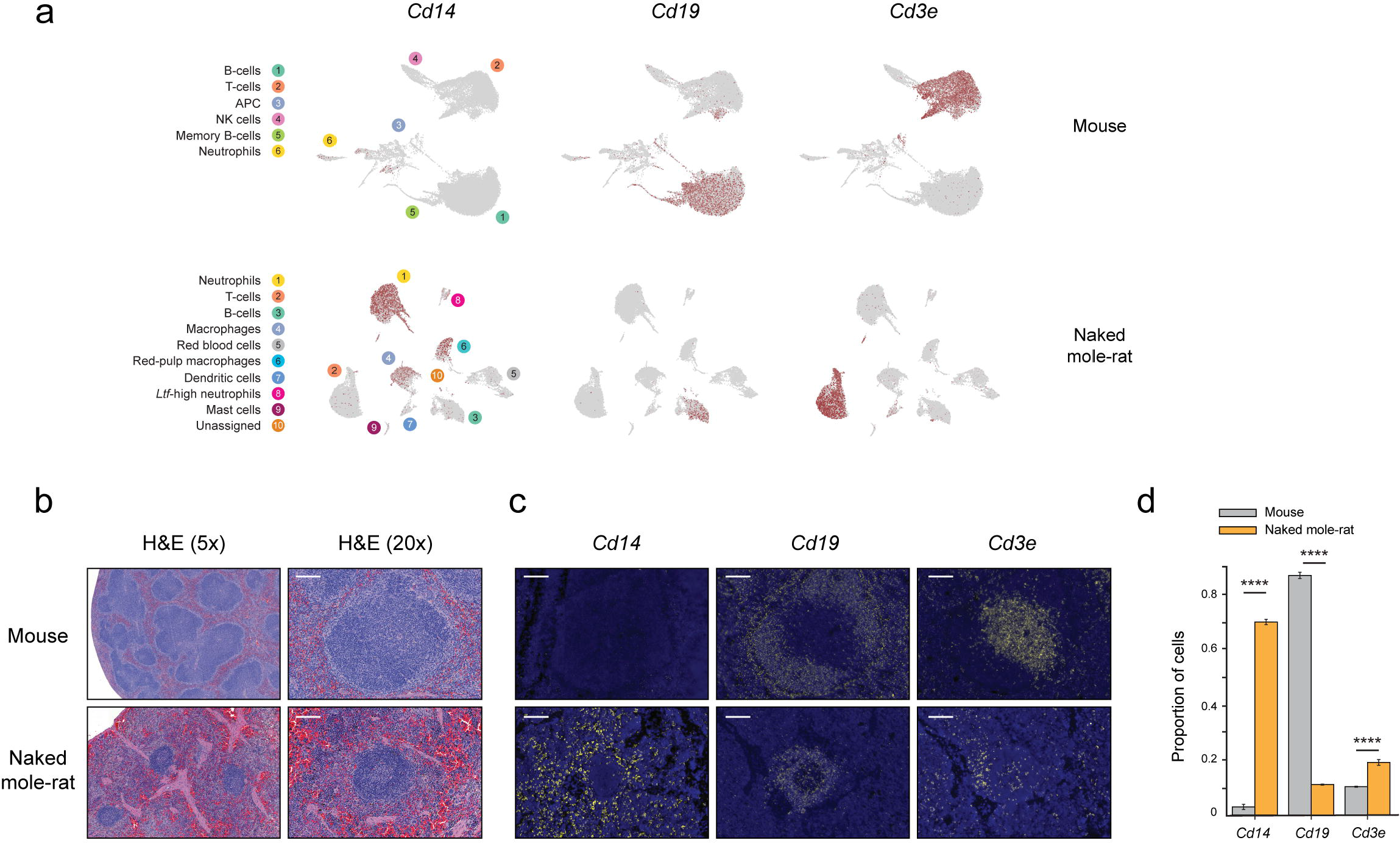
Single-cell RNA sequencing accurately represents the observed microarchitecture of the mouse and naked mole-rat spleens. **a** - Mouse (upper panels) and naked mole-rat (lower panels) spleen cell clusters visualized by UMAP showing the relative expression of *Cd14*, *Cd19* and *Cd3e*. **b** - Representative images of hematoxylin and eosin (H&E) stained sections of mouse (upper panel) and naked mole-rat (lower panel) spleen shown at 5x (left panels) and 20x magnification (right panels). Scale bar = 100μm. **c** - Representative sections of mouse (upper panel) and naked mole-rat (lower panel) spleen showing results from in-situ hybridization staining for *Cd14*, *Cd19* and *Cd3e*. Positive expression of the listed gene was visualized with a TRITC filter (yellow). Nuclear stain was performed using DAPI (blue). Magnification 20x. Scale bar = 100μm. **d** - Quantification of positive staining for *Cd14*, *Cd19* and *Cd3e* in mouse (grey bars) and naked mole-rat (orange bars). Graph shows the proportion (upper panel) of positive staining cells (n=4 biological replicates) and error bars represent the uncertainty associated with the number of puncta which is used as a cutoff to define a cell as positive for a gene (see *Methods*).

We next analyzed the relative distribution and abundance of *Cd14*, *Cd19* and *Cd3e* in the spleens of mice and naked mole-rats. The expression of the housekeeping gene *Rpl13a* and the bacterial-specific *dapB* gene were included as positive and negative controls respectively (see *Methods*). Consistent with our findings from the scRNA-seq data, ISH showed a significantly higher level of *Cd14* in the naked mole-rat compared to the mouse, with staining restricted to the red-pulp regions in both species (Figures 2c-d and Table S5). However, the naked mole-rat has significantly lower abundance of *Cd19*, with positive staining cells localized to the follicular regions of the white pulp (Figures 2c-d and Table S5). Compared with the dramatic differences in the abundance of *Cd14* and *Cd19*, the abundance of *Cd3e* is similar between the two species, which is in line with our scRNA-seq data. Despite their similar relative abundance, the distribution of *Cd3e* positive staining cells is markedly different between the two species, with *Cd3e* expression in the mouse limited to the PALS while that of the naked mole-rat is stochastically distributed throughout both the red and white pulp regions with only minimal aggregation in the PALS (Figures 2c-d). Thus, direct examination of splenic tissue from mouse and naked mole-rat highlights major differences in microanatomy between the two species and supports our results from single-cell sequencing with respect to the proportions of the major immune cell lineages in the two species.

### The naked mole-rat spleen contains nucleated embryonic-like erythroid cells

In addition to the immune cells identified in the naked mole-rat spleen, ~5% of the sequenced cells showed high expression of *hemoglobin* genes but low-expression of the pan-immune marker *Cd45* (*Ptprc*), consistent with their identity as erythroid-lineage (red blood cell-like) cells (cluster 5 in Figure 1f). At convergence, cluster 5 diverged into three sub-clusters (sub-clusters 2, 9 and 10 in Figure S2b) in which we identified variable expression levels of the *ε*-, *γ*- and *β-globin* genes of the *β-globin* locus and the *ζ*-, *α*- and *θ*- globin genes of the *α-globin* locus. Expression of the adult *β*- and *α*- adult globin genes was found to be high in all three sub-clusters, yet not isolated to them (Figures S3a-b respectively), consistent with the capture of cell-free RNA from red blood cells following their lysis (see *Methods*). By contrast, two of the three sub-clusters (sub-clusters 2 and 9) showed high and isolated expression of the *ε*-, *ζ*- and *θ-globin* genes, characteristic of embryonic erythroid cells (16,17) (Figures S3c-e respectively). The fetal *γ-globin* gene showed low expression throughout all clusters (Figure S3f). Furthermore, the *glycophorin c* gene (*Gypc*) and the gene encoding the erythrocyte membrane protein band 4.1 (*Epb41*), whose protein products interact for maintaining erythrocyte shape, deformability and stability (18), showed isolated and correlated expression in sub-clusters 2 and 9 (Figures S4g-j). Similarly, the expression of the gene encoding the erythropoietin receptor (*Epor*), and the transcriptional target of its activation by erythropoietin - the antiapoptotic *Bcl-x(L)* gene, which protects embryonic erythroid cells from apoptosis (19), also showed isolated and correlated expression in sub-clusters 2 and 9 (Figures S3k-n). Finally, the numbers of genes sequenced in sub-clusters 2, 9 and 10 was found to be comparable to that of other myeloid lineage cells, suggesting that they are nucleated (Figure S3o and Table S6). These results indicate that, unlike mice, naked mole-rats have nucleated embryonic forms of erythroid cells in their spleen.

### The naked mole-rat spleen lacks a transcriptionally-defined cell subset that corresponds to natural killer cells in mice

Although lymphoid lineage cells were clearly detected in the naked mole-rat spleen (B-cells marked by high expression of *Cd79a* in cluster 3 and T-cells marked by high expression of *Cd3e* in cluster 2), surprisingly, a cluster corresponding to NK cells was not identified (Figure 1f). In mice, NK cells are transcriptionally defined by high expression of several genes including *Ncr1*, *Gzma*, *Ccl5* and *Nkg7* (Figure 1g and Table S3). Although the naked mole-rat genome assembly and annotation that we used contains an intact *Ncr1* gene (see *Methods*), we did not detect expression of this gene in any naked mole-rat cell profiled by single-cell sequencing in this study. Neither was *Ncr1* detected by RNA sequencing of whole spleen tissue from naked mole-rats. By contrast, we did detect expression of *Gzma*, *Ccl5* and *Nkg7*. All three of these genes were found to be highly expressed in a sub-cluster of the T-cell cluster (cluster 2, Figures 1i and S4a). However, this same sub-cluster of cells also has high expression of both *Cd3e* and *Cd8a*, consistent with their identity as CD8 T-cells, and not NK cells, which lack these T-cell-specific markers in mice and other mammalian species (20) (Figure 1j). Similarly, cluster 9 was found to have high expression of both *Gzma* and *Nkg7,* but also of *Cpa3*, *Tpsab1, Fcer1a* and *Mcpt4*, consistent with their identity as mast cells, and not NK cells (21) (Figures 1k and S4a).

### Genome annotation deficits are not likely a cause of missing natural killer cells in the naked mole-rat

One possibility is that the lack of an NK cell cluster in the naked mole-rat spleen is an artifact that arises due to incomplete annotation of NK cell marker genes in the naked mole-rat genome (see *Methods*). If true, this would mean that reads generated from these marker genes would be filtered as a result of not mapping back to the annotated genome. To test this hypothesis, we simulated a lack of NK cell-marker genes in the mouse by selectively eliminating the marker genes of the mouse NK cell cluster (51 genes, shown in Table S3) and subsequently re-clustered the data. This approach did not change the number of clusters and resulted in only a negligible change with respect to the cellular composition of each of those clusters (Figures S4b-c and Table S7). Thus, even with the absence of NK cell marker genes in the reference mouse transcriptome, we were still able to identify a clear NK cell cluster in the scRNA-seq data from the mouse spleen. This suggests that if the naked mole-rat genome assembly were incompletely annotated for NK cell specific marker genes, it is unlikely to result in the absence of an NK cell cluster in the naked mole-rat spleen dataset.

### Naked mole-rats lack a transcriptionally defined cell subset that corresponds to natural killer cells in circulating immune cells

To investigate the possibility that NK cells are present in the blood, but not in the spleen, we prepared single-cell suspensions of circulating immune cells from the blood of four C57BL/6 mice and four naked mole-rats and determined the gene expression in each sample at single-cell resolution. We captured a total of 12,586 and 10,532 cells from the mouse and naked mole-rat respectively, with a comparable number of genes and UMIs detected per cell to that detected in the spleen (Figures S5a-d). Due to microfluidic failure in the droplet-based sequencing protocol, cells were not captured from one naked mole-rat. We clustered these data similarly to the spleen data, which resulted in a total of six clusters in the mouse (breaking down to a total of nine at convergence) and nine clusters in the naked mole-rat (breaking down to 13 at convergence) (Figures S5e-h and Tables S8-9 respectively).

Mirroring the findings from the spleen, lymphoid cells predominate in the mouse blood (86%) but are less frequent in the naked mole-rat (52%) (Figures S5e-f). B-cells (61%) are the most abundant cell type identified in the mouse whereas neutrophils (45%) are the most common cell type detected in the naked mole-rat blood. A cluster corresponding to NK cells (~4%) was identified in the mouse, but again, not in the naked mole-rat. In an analogous analysis to that undertaken for the spleen, we eliminated the marker genes of the mouse NK cell cluster in circulating immune cells and re-clustered the data. Although the first iteration of our clustering approach eliminated the NK cell cluster by merging NK cells with T-cells (Figure S6a and Table S10), at convergence the NK cell cluster was resolved (Figure S6b and Table S11). Thus, transcriptional profiling of single cells from seven genetically distinct animals, across two organ systems, representing >40,000 total cells failed to identify a transcriptionally distinctive cluster of cells corresponding to NK cells in the naked mole-rat.

### Naked mole-rats lack an expanded major histocompatibility complex class I receptor gene family

Natural killer cells are controlled by a host of variable, germline encoded receptors that recognize major histocompatibility complex class I (MHC-I) molecules (22). In mammals, these MHC-I receptors are encoded in two distinct gene complexes: the *Leukocyte Receptor Complex* (*LRC*), which contains predominantly immunoglobulin-like receptors such as the *killer-cell immunoglobulin-like receptors* (*KIR*) and the *Natural Killer Cell Receptor Complex* (*NKC*), which contains predominantly lectin-like receptors such as *Ly49* (*Klra*), *Cd94* (*Klrd*) and *Nkg*2 (*Klrc*) (23). Both complexes vary in gene content among species. Given the apparent lack of NK cells in the naked mole-rat, we next used a combination of the annotated naked mole-rat genome (24) and a sequence homology search strategy (see *Methods* and Table S12) to generate a complete map of these genomic regions in the naked mole-rat, comparing our results with three well-characterized species and the closely-related guinea-pig (Figures 3a-b).

**Figure 3:**
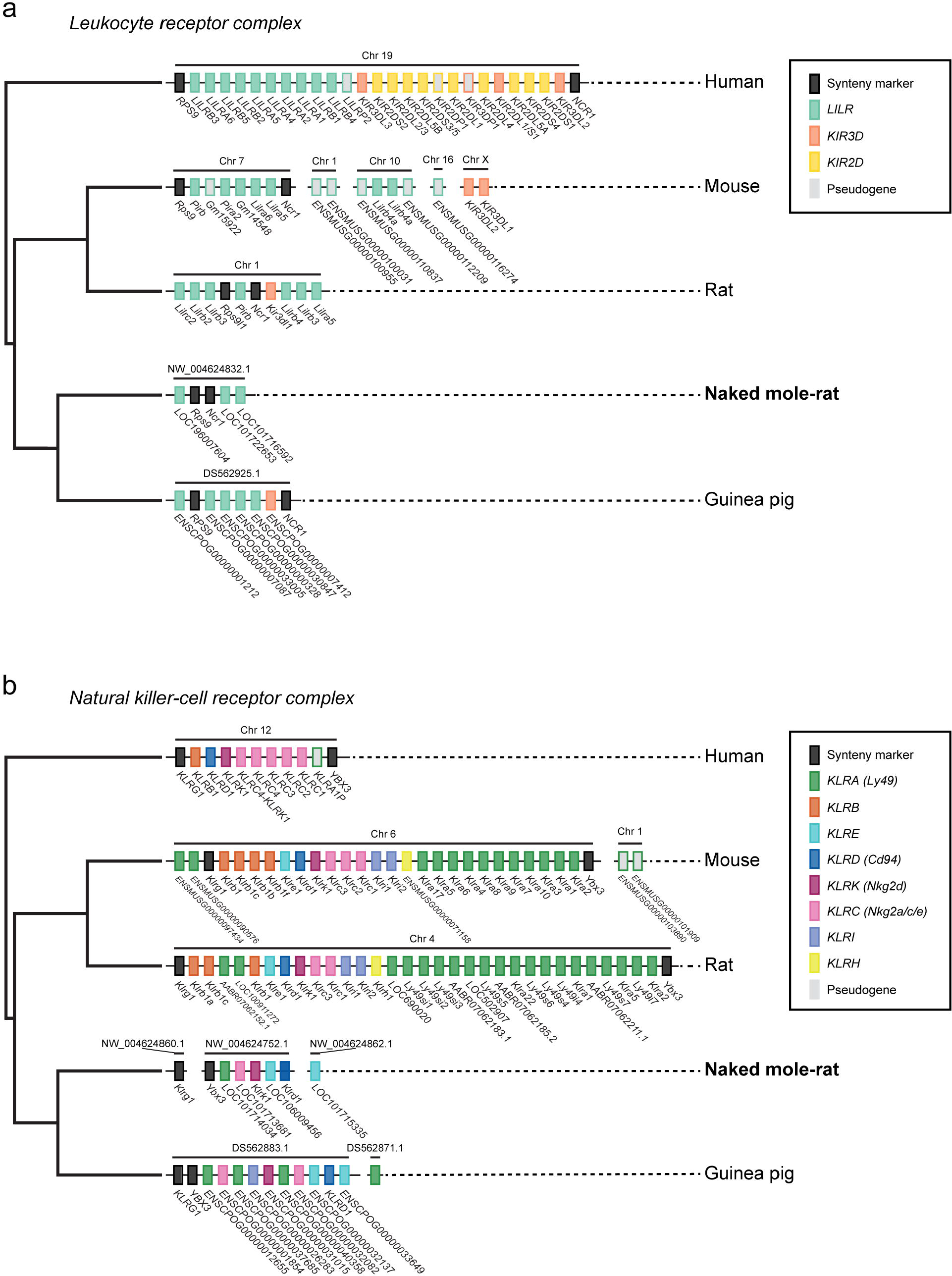
Naked mole-rats lack an expanded family of genes that control natural killer cell responses in other species. Schematic diagram showing the linear arrangement of genes, in 5’-3’ genomic orientation relative to the human genome, in the leukocyte receptor complex (*LRC*) (**a**) and the natural killer cell receptor complex (*NKC*) (**b**) of the human, mouse, rat, naked mole-rat and guinea pig genomes (see also Table S12). Boxes represent genes that are known to control NK cell function, color-coded by family. Synteny markers (black boxes), are the genes that flank the human *LRC* and *NKC*.

Consistent with what is known in the mouse and the rat, we did not find any *KIR* genes in the putative naked mole-rat *LRC* (23). This contrasts with the *LRC* of humans and other primates in which the *KIR* gene family has expanded (25) (Figure 3a). Surprisingly, and in stark contrast to the mouse and rat, which have undergone a dramatic expansion of the *Ly49* gene family in the *NKC* (26), the naked mole-rat genome has only a single inhibitory *Ly49* protein coding gene (a *Klra1* homolog - *LOC101714034*) (Figure 3b), predicted to encode a classical MHC-I receptor (Figure S7a). Our spleen scRNA-seq data revealed that the naked mole-rat *Klra1* homolog is expressed at a low-level, mainly in activated CD8 T-cells (Figure S8a), whereas the mouse *Klra1* is expressed in activated CD8 T-cells, as well as in NK cells (Figure S8b). The *Cd94* and *Nkg2a* genes encode proteins which form an inhibitory heterodimer that recognizes non-classical MHC-I (e.g., HLA-E in humans). Similar to all other compared genomes, a single *Cd94* gene homolog was detected in the naked mole-rat (*Klrd1*, Figures 3b and S7b). However, unlike all other compared genomes only a single *Nkg*2a gene homolog was detected in the naked mole-rat (*LOC101713681*, Figures 3b and S7c). In our spleen scRNA-seq data the naked mole-rat *Cd94* homolog is predominantly expressed in naïve T-cells, activated CD8 T-cells, as well as in red-pulp macrophages (Figure S8a), whereas in the mouse, *Cd94* shows broader expression including in naïve and activated T-cells, macrophages and NK cells (Figure S8b). The naked mole-rat *Nkg2a* homolog was neither detected in the spleen single-cell nor in the whole-tissue RNA-seq data, unlike the mouse *Nkg2a*, which, similar to *Cd94*, shows broad expression across different cell types, including high expression in NK cells (Figure S8b). No other gene family expansions were detected in the putative naked mole-rat *LRC* or *NKC* (Figures 3a-b respectively).

The architecture of the MHC-I receptor gene families in the guinea pig (*Cavia porcellus*) appears to be broadly similar to that of the naked mole-rat (i.e., no expansions of any of the MHC-I receptor gene families in either the *LRC* or the *NKC*, Figures 3a-b). This species, along with the capybara (*Hydrochoerus hydrochaeris*) (which are both members of the Ctenohystrica rodent lineage to which the naked mole-rat belongs), have a population of atypical mononuclear cells with natural killer-like activity called Foa-Kurloff cells (27,28). Most abundant during pregnancy, Foa-Kurloff cells are characterized by the presence of an eccentric reniform nucleus with a single periodic acid-Schiff (PAS)-positive intracytoplasmic inclusion. We examined PAS-stained sections from four healthy two-year old naked mole-rat spleens (two males, two females) as well as routine peripheral blood smears but did not detect a similar population of morphologically distinctive cells in any sample (Figures S9a-b). Taken together, our results show that the naked mole-rat genome lacks a diverse family of MHC-I receptors that control NK cell function in other mammalian species. Further, we show that this unusual feature may be more widespread in mammalian genomes than previously recognized.

### In vivo response to lipopolysaccharide challenge

Our data suggest that the naked mole-rat immune system is characterized by a predominance of myeloid lineage, innate immune cells with high expression of *Cd14*. *Cd14* encodes a protein that functions as a co-receptor with Toll-like receptor 4 for the detection of bacterial lipopolysaccharide (LPS) and other pathogen-associated molecular patterns (29,30). In order to examine how the naked mole-rat immune system responds to an infection-mimicking challenge, we profiled the transcriptional response to administration of LPS.

Four mice and four naked mole-rats were randomly distributed into either a control or treatment group (two animals from each species per group). Animals assigned to the control group received sterile saline by intraperitoneal injection. Animals assigned to the treatment group received 1mg/kg bodyweight LPS by intraperitoneal injection (Figures 4a-b). Four hours following administration of either saline or LPS, animals were sacrificed, and the transcriptional responses in circulating and spleen-resident immune cells in each animal were assessed at whole tissue and single-cell resolution.

**Figure 4:**
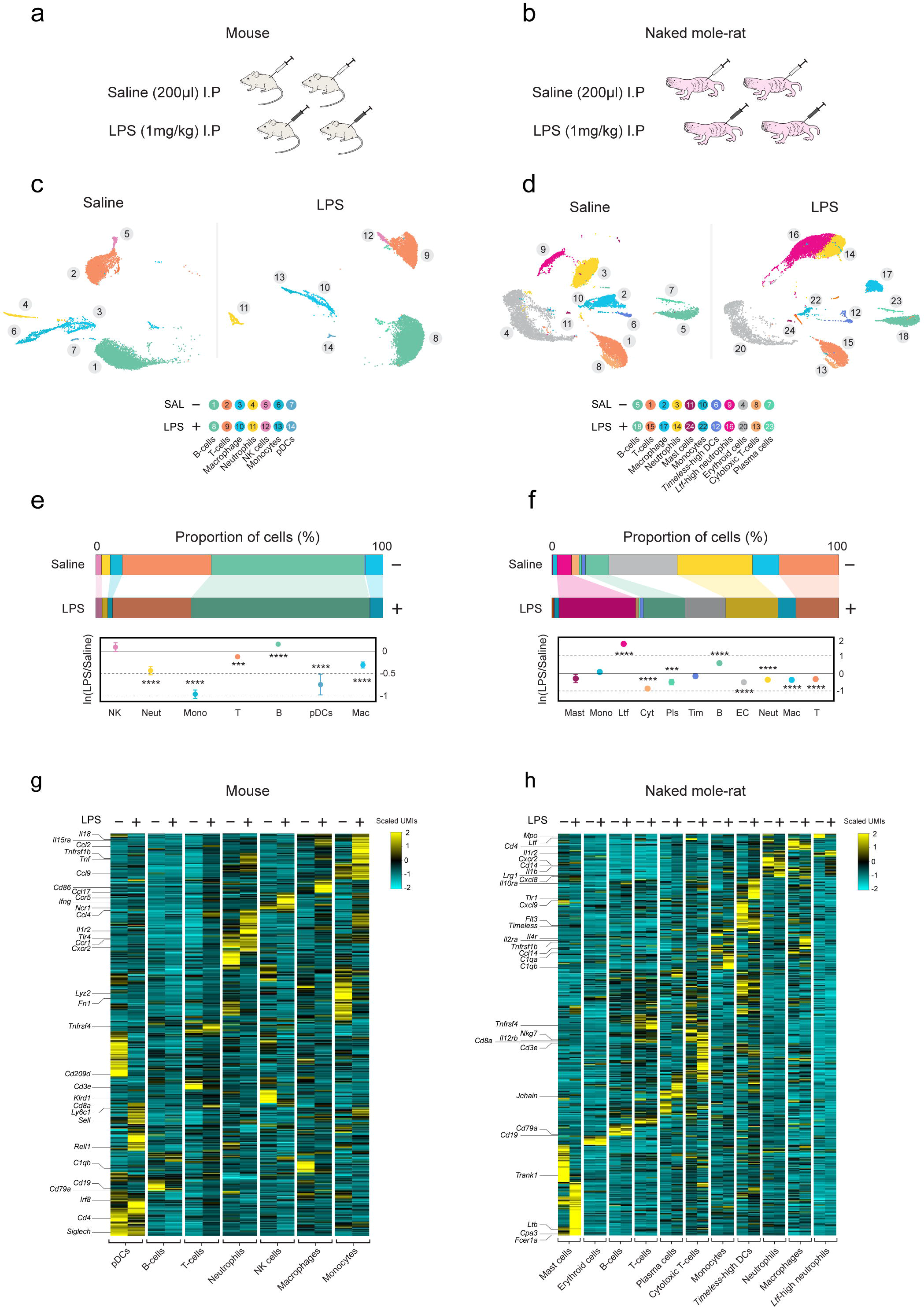
Single-cell transcriptional profiling reveals that naked mole-rats have a lipopolysaccharide-responsive cell subset not found in mice. **a** - Schematic showing the experimental outline in which four C57BL/6 mice are assigned to either saline control (n=2) or LPS-challenged (n=2) groups. After four hours, single-cell suspensions were derived from the spleen of each animal and subjected to single-cell RNA sequencing. **b** - Schematic showing the experimental outline in which four naked mole-rats are assigned to either saline control (n=2) or LPS-challenged (n=2) groups. After four hours, single-cell suspensions were derived from the spleen of each animal and subjected to single-cell RNA sequencing. **c** - Mouse spleen cell clusters from saline control (left panel) and LPS-challenged (right panel) visualized by UMAP. Colors indicate clusters that correspond to the listed cell types. **d** - Naked mole-rat spleen cell clusters from saline control (left panel) and LPS-challenged (right panel) visualized by UMAP. Colors indicate clusters that correspond to the listed cell types. **e** - Stacked bar charts showing the relative proportions (%) of the listed cell types in saline control and LPS-challenged mice (upper panel) with the ln(LPS/saline) cell-count ratios are shown as effect sizes (lower panel). **f** - Stacked bar charts showing the relative proportions (%) of the listed cell types in saline control and LPS-challenged naked mole-rats (upper panel) with the ln(LPS/saline) ratios are shown as effect sizes (lower panel). **g** - Heatmap showing the gene expression in mouse spleen cells from saline control (−) and LPS-challenged (+) animals in each of seven clusters (mean across the cells of each cluster) that correspond to the listed cell types. Selected marker genes are listed to the left. **h** - Heatmap showing the gene expression in naked mole-rat spleen cells from saline control (−) and LPS-challenged (+) animals in each of eleven clusters (mean across the cells of each cluster) that correspond to the listed cell type. Selected marker genes are listed to the left.

At the whole tissue level, RNA sequencing data revealed significant changes in gene expression following LPS challenge in both species (Figures S10a-d and Tables S13-14), evidenced by upregulation of inflammatory mediators including *Il1a*, *Il6*, *Il10*, *Il15*, *Ifng* and *Tnf*. Gene set enrichment analyses showed broadly similar changes between the two species with activation of the inflammatory response via the *NF-KB* pathway highlighted as a prominent feature (Figures S10e-f and Tables S15-16).

### Single-cell RNA-sequencing reveals divergent species responses following lipopolysaccharide challenge

Single-cell data revealed that spleen cells from both the control and LPS-challenged animals of both species were broadly classified into similar cell types to those observed in untreated animals (Figures 1e-f and 4c-d). However, we observed significant changes in the relative proportion of cells between control and LPS-challenged animals. In the mouse we observed marginal increases in the relative proportion of NK cells and B-cells, and decreases in the relative proportion of neutrophils, monocytes, T-cells, macrophages and dendritic cells (Figure 4e). We observed similar changes in the relative proportion of most cell types in the naked mole-rat, with the exception of *Ltf*-high neutrophils, which showed a dramatic five-fold increase (from ~5% in control animals to >25%) following the LPS challenge (adjusted p-value < 10^−22^) (Figure 4f).

Marked transcriptional responses were evident following LPS challenge for each cell type compared between the control and treatment groups in each of the species (Figures 4g-h). As described previously by Shalek et al. (31), these LPS-responsive transcriptional changes were driven by *expression change* (a change in the level of the expression among the cells which expressed a given gene in both conditions) and/or *expression-induction change* (a change in the fraction of cells expressing a certain gene), illustrated in our data by the transcriptional changes of *Nfkbia* (Figures 5a-b). We quantified these expression and expression-induction effects for each gene across each of the cell types in both species using a hurdle model (see *Methods*) (Figures 5c-d). This revealed that, qualitatively, splenic monocytes and neutrophils experience the strongest difference in expression changes between the two species (Figure 5c). For expression changes, only mouse NK cells showed statistically significant enrichments (adjusted p-value < 0.1) of the *Ifnα*, *Ifnγ* and the *NF-KB* pathways (Figure 5e and Table S17). By contrast, all cell types in the mouse, with the exception of NK cells, showed statistically significant expression-induction enrichments (adjusted p-value < 0.1), predominantly in the *Ifnα*, *Ifnγ* and the *NF-KB* pathways (Figure 5f and Table S18). In the naked mole-rat, we only observed statistically significant enrichments (adjusted p-value < 0.1) for expression-induction changes mostly in the *Ifnα* and *Ifnγ* pathways in splenic monocytes (Figure 5g and Table S19). These results thus indicate that, for both species, many of the transcriptional responses to LPS challenge observed at the whole tissue level are contributed by expression-induction changes at the single-cell level.

**Figure 5:**
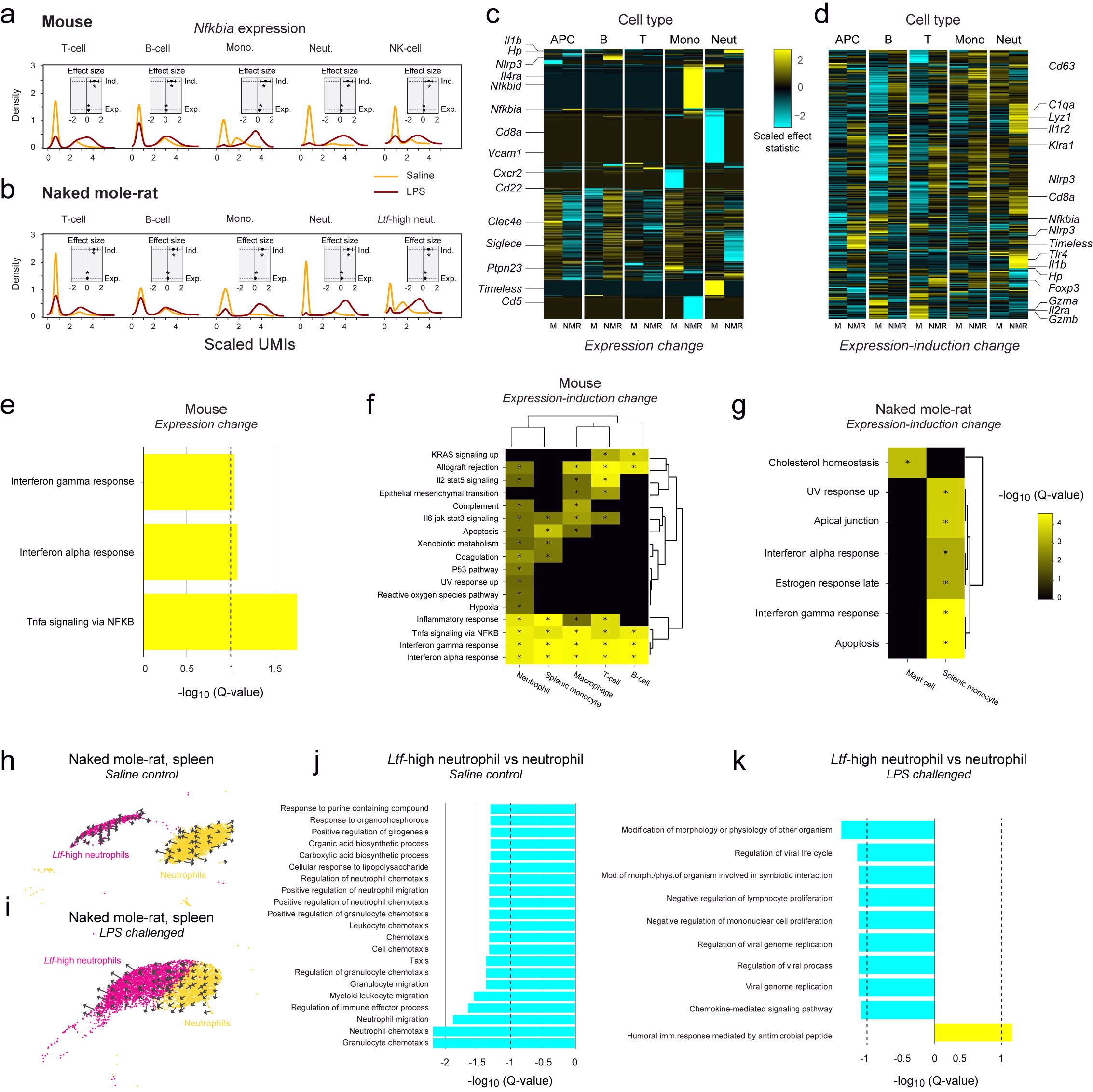
Cell specific responses to lipopolysaccharide challenge in the naked mole-rat and mouse spleen. **a** - Selected density plots showing the expression distribution of *Nfkbia* from saline control (orange) and LPS-challenged (red) mice across the cells of five representative cell types. Insets show the estimated treatment expression (Exp.) and expression-induction (Ind.) effect sizes (LPS relative to saline) (asterisks mark adjusted p-value < 0.05). **b** - Selected density plots showing the expression distribution of *Nfkbia* from saline control (orange) and LPS-challenged (red) naked mole-rats across the cells of five representative cell types. Insets show the estimated treatment expression (Exp.) and expression-induction (Ind.) effect sizes (LPS relative to saline) (asterisks mark adjusted p-value < 0.05). **c** - Heatmap showing the estimated treatment-effect statistics for expression change. Selected marker genes are shown to the left. **d** - Heatmap showing the estimated treatment-effect statistics for expression-induction change. Selected marker genes are shown to the right. **e** - Bar chart showing the gene set enrichment analyses (GSEAs) of the intra-species treatment effect on expression change in mouse NK cells. The x-axis reports the log_10_ adjusted p-value (q-value) of the GSEA. The log_10_ adjusted p-value is signed and color-coded by the direction of the effect (up- and down -regulation in LPS relative to saline: yellow and cyan respectively). Black vertical dashed lines correspond to an adjusted p-value of 0.1. **f** - Heatmap showing GSEAs of the intra-species treatment effects on expression-induction change in mice across the listed cell types. The log_10_ adjusted p-value (q-value) of the GSEA is indicated by the shade of the color and the colors encode the direction of the effect (up- and down -regulation in LPS relative to saline: yellow and cyan respectively). Gene sets with an adjusted p-value of < 0.1 are indicated with an asterisk. **g** - Heatmap showing GSEAs of the intra-species treatment effects on expression-induction change in naked mole-rats across the listed cell types. The log_10_ adjusted p-value (q-value) of the GSEA corresponds is indicated by the shade of the color and the colors encode the direction of the effect (up- and down -regulation in LPS relative to saline: yellow and cyan respectively). Gene sets with an adjusted p-value of < 0.1 are indicated with an asterisk. **h** - Clusters representing *Ltf*-high neutrophils (pink) and neutrophils (yellow) from saline control animals visualized by UMAP. Overlaid on each cluster are arrows that represent the cell trajectory as inferred by RNA velocity. **i** - Clusters representing *Ltf*-high neutrophils (pink) and neutrophils (yellow) from LPS-challenged animals visualized by UMAP. Overlaid on each cluster are arrows that represent the cell trajectory as inferred by RNA velocity. **j** - Bar chart showing gene set enrichment analyses (GSEAs) comparing *Ltf*-high neutrophils to neutrophils from saline control naked mole-rats. The x-axis reports the log_10_ adjusted p-value (q-value) of the GSEA. The log_10_ adjusted p-value is signed and color-coded by the direction of the effect (up- and down -regulation in *Ltf*-high neutrophils relative to neutrophils: yellow and cyan respectively). Black vertical dashed lines correspond to an adjusted p-value of 0.1. **k** - Bar chart showing GSEAs comparing *Ltf*-high neutrophils to neutrophils from LPS-challenged naked mole-rats. The x-axis reports the log_10_ adjusted p-value (q-value) of the GSEA. The log_10_ adjusted p-value is signed and color-coded by the direction of the effect (up- and down - regulation in *Ltf*-high neutrophils relative to neutrophils: yellow and cyan respectively). Black vertical dashed lines correspond to an adjusted p-value of 0.1.

### Lipopolysaccharide challenge induces transition of neutrophils to Ltf-high neutrophils in the naked mole-rat spleen

Accompanying the increase in the relative proportion of *Ltf*-high neutrophils following LPS challenge, the UMAP embedding of LPS-challenged and control cells from the naked mole-rat spleen showed a convergence between the neutrophil and *Ltf*-high neutrophil clusters (Figures 5h-i). This result suggests a transition, from neutrophils under control (saline vehicle) conditions to *Ltf*-high neutrophils under LPS challenge conditions. In order to investigate this convergence of cell types, we used *RNA velocity* (32) to estimate the trajectories of the cells in the neutrophil and *Ltf*-high neutrophil clusters under control and LPS challenge conditions. Results from this analysis indicate that while there is no trajectory between neutrophils and *Ltf*-high neutrophils under control conditions (Figure 5h), LPS challenge appears to induce a trajectory from neutrophils towards *Ltf*-high neutrophils (Figure 5i), suggesting that exposure to LPS induces neutrophils to transition into *Ltf*-high neutrophils. Supporting this conclusion is the significant increase in the relative proportion of *Ltf*-high neutrophils and decrease in the relative proportion of neutrophils (adjusted p-values < 10^−22^) following LPS challenge (Figure 4f).

We next characterized the gene-expression differences between *Ltf*-high neutrophils and neutrophils at the biological pathway level (see *Methods*). Only pathways significantly enriched in genes downregulated in *Ltf*-high neutrophils relative to neutrophils were found in cells from the control condition (adjusted p-value < 0.1). These pathways predominantly relate to granulocyte chemotaxis, suggesting that either these pathways are activated in neutrophils or inhibited in *Ltf*-high neutrophils (Figure 5j and Table S20). Following LPS challenge, genes upregulated in *Ltf*-high neutrophils relative to neutrophils showed enrichment of humoral immune responses mediated by antimicrobial peptides, whereas downregulated genes showed enrichment of viral replication processes, host-pathogen interactions and negative regulation of immune cell proliferation (adjusted p-value < 0.1) (Figure 5k and Table S21). These findings are in accordance with the observations of the changes in cell-type relative proportions and trajectories, indicating a transition of neutrophils to antimicrobial *Ltf*-high neutrophils upon LPS challenge.

### Response to lipopolysaccharide challenge in circulating immune cells

In order to determine the systemic immune response to infection, we also performed scRNA-seq on circulating immune cells in the blood from each mouse and naked mole-rat in the saline control and LPS challenged groups (Figures S11a-h). This analysis revealed the same types of transcriptional responses to LPS challenge in cells compared between the two conditions in both species (Figures S12a-b). We thus fitted the hurdle model to these data (Figures S12c-d). Mirroring our findings from the spleen, for expression changes, only the mouse NK cells showed statistically significant (adjusted p-value < 0.1) enrichments of the *Ifnα*, *Ifnγ* and the *NF-KB* pathways (Figure S12e and Table S22). For expression-induction changes, all mouse cell types with the exception of NK cells showed statistically significant (adjusted p-value < 0.1) enrichments, predominantly in the *Ifnα*, *Ifnγ* and the *NF-KB* pathways (Figure S12f and Table S23). In the naked mole-rat, we only observed statistically significant enrichments (adjusted p-value < 0.1) for expression-induction changes. Specifically, B-cells showed most of the enrichments in the *Ifnα*, *Ifnγ*, and *NF-KB* pathways, whereas among other pathways, T-cells also showed enrichment for the complement system pathway (Figure S12g and Table S24).

The increase in the relative proportion of *Ltf*-high neutrophils in the circulating immune cell pool was even more pronounced than that observed in the spleen, increasing from ~0.5% in control animals to >56% in response to LPS challenge (adjusted p-value < 10^−22^) (Figure S11f). This dramatic increase in systemic *Ltf*-high neutrophils was accompanied by a larger decrease in the relative proportion of neutrophils (from ~78% to ~14%) (adjusted p-value < 10^−22^), again suggesting that LPS challenge induces transition of neutrophils to *Ltf*-high neutrophils. Notwithstanding, the changes in cell trajectories as evidenced by the RNA velocity analysis were less clear than those identified in the spleen (Figures S12h-i). Under saline control conditions, differentially expressed genes were not found to be enriched in any specific biological pathway (adjusted p-value < 0.1) when comparing *Ltf*-high neutrophils to neutrophils. By contrast, and in accordance with our observations from the spleen, the genes upregulated in *Ltf*-high neutrophils relative to neutrophils following LPS challenge were found to be enriched in antimicrobial immune responses, and downregulated genes were found to be enriched in modification of host-pathogen interactions (adjusted p value < 0.1) (Figure S12j and Table S25).

Taken together, the results of our LPS challenge experiments show that despite a comparatively conserved transcriptional response between the mouse and naked mole-rat at the whole-tissue level, these species show divergent responses at single-cell resolution, with the naked mole-rat undergoing a significant up-regulation in the number of its enigmatic *Ltf*-high neutrophil population.

## Discussion

Using single-cell gene expression profiling and comparative genomics, we investigated if the naked mole-rat immune system was similar to that of the genetically and functionally well-characterized laboratory mouse. We discovered that the naked mole-rat immune system has an unusual composition, being defined by a higher myeloid to lymphoid cell ratio, a lack of classically defined NK cells, a dearth of germline encoded NK cell receptors and the presence of an LPS responsive neutrophil-like cell population with high expression of *lactotransferrin* and *cathelicidin*. In addition, the naked mole-rat spleen appears to contain a population of nucleated embryonic-like erythroid cells, although this finding deserves further characterization to rule out a potential bias introduced during red blood cell lysis. These data represent the first comprehensive exploration of the naked mole-rat immune system and may provide critical clues for understanding the cancer-resistance and exceptional longevity of this model species.

Most vertebrate species have a population of innate lymphocytes that recognize and remove virally infected or malignantly transformed cells (33). These effector functions rely on a suite of germline encoded cell-surface receptors that discriminate subtle differences in the expression level of MHC-I molecules on target cells (34). In a classic example of convergent evolution, divergent mammalian species have expanded separate gene families that encode MHC-I receptors with almost identical function. Illustrating this phenomenon, the primate lineage has expanded the immunoglobulin-like *KIR* genes, resulting in 14 distinct genes in the human whereas the mouse and the rat have expanded the lectin-like *Ly49* (*Klra*) receptors, with over 30 distinct genes now described in these two species alone (26). More recently, the expansion of yet another lectin-like MHC-I receptor gene family, *Cd94/Nkg2a*, has been described in a number of bat species (35). Contrasting with these examples, the naked mole-rat lacks both a cell population that approximates the transcriptional signature of NK cells described in other species and also lacks an expanded family of lectin – or immunoglobulin-like MHC-I receptors that represent their prototypical control mechanisms (20).

Structural or allelic polymorphism in NK cell receptor gene families is a common feature of other mammalian species and are thought to provide a selective advantage by increasing the genetic (and hence functional) diversity of this arm in the immune system (22). That we did not collect population-level genomic data in naked mole-rats raises the possibility that we are underestimating the diversity of these MHC-I receptor gene families in this species. Notwithstanding this limitation, the lack of an expanded family of MHC-I receptors in the naked mole-rat reference genome is still most unusual amongst mammals, although isolated examples, such as the marine carnivores, have been described (36). However, that our study revealed a similar architecture in the guinea pig genome, raises the possibility that this genomic state is ancestral and may therefore be more common among mammalian genomes than previously thought.

Our data show that the naked mole-rat has a higher proportion of myeloid cells, abundance of *Cd14* expression, and a seemingly unique population of LPS-responsive neutrophil-like cells with high expression of genes that encode anti-bacterial peptides. Further, although the naked mole-rat genome is not replete with an unusually large number of genes that encode antimicrobial peptide products, those that have been studied show potent antimicrobial activity (37). Taken together, these findings suggest that the naked mole-rat immune system has evolved under stronger anti-bacterial than anti-viral immune selective pressure. Supporting this assertion, although individually limited in their scope, there is evidence that the naked mole-rat is unusually susceptible to viral infection (38,39). In addition, the xenophobic eusocial structure of naked mole-rats, combined with their strictly subterranean ecological niche, suggests that viral selective pressure on naked mole-rats might be weaker than in other mammalian species such as bats, in which intricate anti-viral defenses have been described (35).

Taken together, our study adds further support to the notion that, although seemingly common, an expanded family of MHC-I receptor genes is not a prerequisite for the proper functioning of innate immunity in mammals. A more widespread evolutionary investigation is likely to be informative in determining the frequency of this genomic architecture across mammalian genomes and will likely shed light on the extent to which the naked mole-rat is an outlier in this regard. Similarly, an investigation into the frequency of species that lack classical NK cells is likely to have important bearings on the role of innate lymphocyte-mediated immunity in mammals.

## Methods

### Animals

In this study we used comparative histomorphology, comparative genomics, fluorescent in-situ hybridization and deep transcriptional profiling to study the immune systems of mice and naked mole-rats. The animals used in this study comprised 12 eight to ten-week-old C57BL/6 mice (six males, six females) and 12 23-25 month-old naked mole-rats (six males, six females). The naked mole-rats were part of a well-characterized colony (1) and the mice were purchased from the Jackson Laboratories (Bar Harbor, Maine, USA) and maintained in the vivarium for at least two weeks prior to use. The ages selected yielded young, healthy individuals that were physiologically age-matched (~5-10% of observed maximum lifespan). Both species were maintained on a 12h dark-light cycle and provided food ad libitum. Mice were provided water ad libitum. In accordance with standard colony management, naked mole-rats did not receive supplemented water as the water content of their diet is sufficient for maintaining appropriate hydration. All animal use and experiments were approved by the Buck Institute Institutional Animal Care and Use Committee (IACUC protocol number A10173).

### Organ collection and processing

We investigated the immune-cell repertoire of eight naked mole-rats (four males, four females) and eight C57BL/6 mice (four males, four females) using single-cell transcriptional profiling of the spleen and circulating immune cells. Organ collection was performed on all animals between 8am and 10am. Animals were anesthetized using isoflurane. Blood was drawn via cardiac puncture and immediately transferred into EDTA-containing tubes, mixed by inversion and placed on ice.

1. *Blood:* One ml of EDTA-treated blood was resuspended in 20ml ACK lysis buffer (Gibco A1049201) for 5 minutes at 21°C. Cells were pelleted (500 × *g*, 10mins, 4°C), resuspended in PBS with 5% FBS and washed twice. Following resuspension, cells were passed through a 40μm filter and placed on ice prior to determination of viability and density. Visual inspection of trypan blue (Gibco 15250061) stained cells loaded on a hemocytometer slide was used to determine the density and viability of cells in the suspension.
2. *Spleen:* Immediately following cardiac puncture the spleen was removed by dissection, transferred to a sterile petri dish containing PBS with 5% FBS and minced using razors. Following trituration with a 5ml serological pipette, the spleen fragments were ground through 100μm and 40μm cell strainers (Falcon 352360 and 352340) with a syringe plunger. Cells were pelleted (500 × *g*, 10mins, 4°C), resuspended in 20ml ACK lysis buffer (Gibco A1049201) for 5 minutes at 21°C and washed twice with PBS with 5% FBS. Following resuspension, cells were passed through a 40μm filter and placed on ice prior to determination of viability and density. Visual inspection of trypan blue (Gibco 15250061) stained cells loaded on a hemocytometer slide was used to determine the density and viability of cells in the suspension.

### Lipopolysaccharide challenge

In order to study the transcriptional response of mice and naked mole-rats to stimulation of the innate immune system we performed transcriptional profiling four hours following challenge with lipopolysaccharide (LPS). Two animals from each species (one male, one female) were randomly assigned into either challenge or control groups (total = four mice, four naked mole-rats). Control animals received intraperitoneal injection of 200μl sterile saline. Challenged animals received intraperitoneal injection of 1mg/kg LPS (LPS-EB VacciGrade, *Escherichia coli* 0111:B4, Invivogen, San Diego), prepared in sterile saline to a final volume of 200μl. Body temperature of all animals was monitored by rectal probe every 30mins for two hours following injection. Collection and processing of blood and spleen was performed four hours following initial injection and as described above with the exception that approximately one fifth of each spleen was immediately flash-frozen in liquid nitrogen (for whole-tissue RNA extraction and sequencing).

### Single-cell RNA sequencing and data analysis

Single cells were captured in droplet emulsion using the Chromium Controller (10X Genomics) and scRNA-seq libraries were constructed according to the 10X Genomics protocol using the Chromium Single-Cell 3’ Gel Bead and Library V2 Kit. In brief, cell suspensions were diluted in PBS with 5% FBS to a final concentration of 1 × 10^6^ cells/ml. (1000 cells per μl). Cells were loaded in each channel with a target output of 3000 cells per sample. All reactions were performed in a Bio-Rad C100 Touch Thermal Cycler with a 96 Deep Well Reaction Module. Twelve cycles were used for cDNA amplification and sample index PCR. Amplified cDNA and final libraries were evaluated using a Bioanalyzer 2100 (Agilent Technologies) with a high sensitivity chip. Samples were sequenced on an Illumina HiSeq 4000.

ScRNA-seq fastq files were demultiplexed to their respective barcodes using the 10X Genomics Cell Ranger mkfastq utility. Unique Molecular Identifier (UMI) counts were generated for each barcode using the Cell Ranger count utility. The mm10 reference genome and the mouse Gencode version M12 annotation were used for mapping the mouse reads (40). The HetGla2.0 reference genome along with the RefSeq GCF_000247695.1 annotation, to which we manually added the annotation of the *Ncr1* gene from Ensembl’s *Heterocephalus glaber* female annotation (which is the only naked mole-rat annotation that covers this gene), were used for mapping the naked mole-rat reads. For each sample, barcodes that were not likely to represent captured cells were filtered out by detecting the first local minimum above 2 in a distribution of log_10_(#UMIs). Similarly, for each sample, genes that are too sparsely captured across barcodes were filtered out by detecting the first local minimum above 3 in a distribution of log_10_(#barcodes). Finally, barcodes capturing more than a single cell (multiplets) were sought as local modes in the distributions of log_10_(#genes) and log_10_(#UMIs), whose x-axis maxima are more than 1.5 higher than that of the x-axis location of the global maximum of the respective distribution and includes less than 5% of the total number of barcodes. In other words, barcodes that, based on the number of genes or UMIs they captured, appear inflated with respect to all other barcodes, were regarded as multiplets and thus filtered (summary statistics shown in Table S26).

To identify cell clusters, the samples from each species were concatenated, UMIs were then scaled to the read-depth of their respective barcodes, and genes with high expression dispersion were obtained using Seurat (41) (Table S26). Principal Components Analysis (PCA) was then performed on these gene × cell scaled UMI matrices using the rsvd (42) R (43) package to reduce the dimensionality of the data, retaining the 50 PCs explaining the highest amount of variation. We then used Seurat’s methodology to build a Shared Nearest Neighbor (SNN) graph of these cell-embedding data, first generating a K-Nearest Neighbor (KNN) graph using K = min(750, #cells-1) and a Jaccard distance cutoff of 1/15. The SNN graph was then used as input to the Louvain algorithm, implemented in the ModularityOptimizer software (44). Since this implementation uses a resolution parameter that strongly affects the number of clusters, we searched the 0.05-1.225 range of this parameter, using the mean unifiability isolability clustering metric as our maximization parameter. This process was initially done on all cells in our data, and subsequently repeated for each cluster individually, in an iterative manner where convergence was defined as not being able to break down a cluster into sub-clusters (see Tables S1-2 and S8-9 for the cell-cluster assignments for each species-tissue data set).

In order to obtain gene markers for each cluster, we ran a differential expression test implemented in Seurat (using a likelihood ratio test between a model that assumes that a gene’s expression values of two compared clusters values were sampled from two distributions, versus a null model which assumes they were sampled from a single distribution). A marker gene of a given cluster was defined as a gene which was found to be significantly overexpressed (multiple-hypotheses adjusted (45) p-value < 0.05) in that cluster compared to all other clusters (see Tables S3-4 for the cluster marker genes for the mouse and naked mole-rat spleen data respectively). Cell-type assignment to the clusters was performed manually using a panel of known marker genes. In all genes × cells heatmap figures, the total number of cells was reduced to a maximum of 2,500 (in proportions with respect to the number of cells per each cluster) by randomly down-sampling cells in each cluster.

### Obtaining a list of mouse–naked-mole-rat orthologous genes

To obtain mouse-naked mole-rat one-to-one gene orthologs we applied a reciprocal-blast approach (46). Specifically, we used BLASTp for each of the naked mole-rat protein sequences against all mouse protein sequences and vice versa, and the same for transcript sequences using BLASTn. In both cases, we retained only hits with e-value < 0.1 and a sequence identity >80%. Any gene for which any of its protein or transcript forms reciprocally hit back any of the protein or transcript forms of that gene was retained as a one-to-one gene ortholog (Table S27).

In order to detect any possible member of any of the MHC-I receptor gene families in the annotations of the naked mole-rat and guinea pig genomes (where for guinea pig we used the Cavpor3.0 - GCA_000151735.1 assembly along with its Ensembl Cavpor3.0 annotation), we used the OrthoFinder software, which finds groups of orthologous genes (47). Specifically, OrthoFinder was used with the human (hg38 genome assembly along with its Gencode v25 annotation (48)), mouse, rat (Rnor_6.0 - GCF_000001895.5 genome assembly along with its Ensembl Rattus_norvegicus.Rnor_6.0 annotation), naked mole-rat and guinea pig protein sequences as input. The resulting orthogroups were then assigned to one of the MHC-I receptor gene families (Figure3 and Table S12) based on the human, mouse, and rat annotations. Each putative functional NK cell receptor gene in the naked mole-rat was individually screened for the presence of three sequence motifs indicative of their likely function: 1. Immunoreceptor tyrosine inhibitory motifs (ITIM), 2. immunoreceptor tyrosine activating motifs (ITAM) and 3. double-cysteine residues.

### Testing for different numbers of sequenced genes across single-cell RNA-sequencing clusters

To test whether the numbers of sequenced genes differ between the different myeloid lineage cells in the naked mole-rat we fitted a negative binomial regression to the gene-count data (to account for over-dispersion), using the MASS (49) R (43) package, where cell-type was specified as a categorical factor, with the naked mole-rat spleen sub-cluster 1 set as baseline.

### Cell-type specific responses to lipopolysaccharide challenge

In single-cell data (including the data generated in this study) the distribution of the number of UMIs per a given gene across the cells in a certain cluster is typically bimodal, with a left mode located at zero and a right mode located at higher than zero. Across different factors (e.g., treatments, different cell types) a change in the fraction of cells expressing the gene, i.e., a change in expression-induction, can be explicitly quantified as well as the change in the levels of expression among the cells expressing the gene (i.e., a shift in the right modes of the distributions of the number of UMIs). To this end, in all our differential expression analyses, with the exception of the one applied for defining cluster marker genes, we fitted a hurdle generalized linear mixed-model to the data, using the lme4 (50) R (43) package. Specifically, effects on expression-induction were estimated by fitting a logistic mixed-effects regression model (with a binomial family using the logit link function) by binarizing the outcome to 0 (not expressed in the cell) and 1 (expressed in the cell). Effects on expression levels were estimated by fitting a mixed-effects regression model (with a *gamma* family using the log link function), only to the cells in which the gene was expressed. In both cases, sample was defined as a random effect.

In order to detect intra-species cell-type-specific responses following LPS challenge, for each gene in each pair of matching cell types in the saline and LPS data sets (Tables S28-29 for the spleen and circulating immune cell datasets respectively), we fitted the hurdle model, described above, to the LPS and saline data. Specifically, to estimate the effect of treatment on induction of expression as well as on change of expression levels, treatment was specified as a fixed categorical effect, with saline set as baseline, and sample was specified as a random effect, as noted above. The heatmaps displaying the expression-changes and expression-induction changes (Figures 5c-d for the spleen data and S12c-d for the circulating immune cells data respectively) are limited to the set one-to-one orthologs gene between mouse and naked mole-rat (Table S27) and to cell types we matched between them (Tables S28-29 for the spleen and blood circulating immune cells datasets respectively).

In order to perform a Gene Set Enrichment Analyses (GSEA), for any of the fitted models, for each cell type, we ranked the genes by the p-value of the estimated effect and used that as an input to the fgsea function in the fgsea (51) R (43) package. We used the Hallmark gene sets of the MSigDB collection (52) for GSEAs with the exception of the *Ltf*-high neutrophil expression-change GSEAs in which we used the Gene-Ontology Biological Pathways sets (51,53). In all GSEAs, we performed multiple hypotheses p-value adjustments (45) for each cell type. Although not as conservative as adjusting the p-values across all cell types, the latter approach would suffer from being imbalanced between the two species for which a different number of cell types was tested. To perform intra- and inter-species comparisons between proportions of cell types we fitted a multinomial regression (similar to the mixed-effects models fitted to the expression data, except for the sample random effect), implemented by the nnet (49) R (43) package.

### Comparative splenic histomorphology and fluorescent in-situ hybridization

We investigated the histomorphology and expression of four genes in the spleen of four naked mole-rats (two males, two females) and four C57BL/6 mice (two males, two females). Organ collection was performed as described above and spleens were immediately transferred into 10% neutral-buffered formalin. Following fixation, all samples were trimmed, routinely processed, embedded and sectioned at 4μm thickness in an RNase-free environment. Consecutive tissue sections were stained with hematoxylin and eosin (H&E), periodic acid-Schiff (PAS) and the expression of *Cd3e*, *Cd19*, *Cd14* and *Rpl13a* was determined by fluorescent in-situ hybridization (ISH) (ViewRNA, Invitrogen). The expression of the bacterial gene *dapB* was determined as a negative control.

Samples were stained with 4’,6’-diamidino-2-phenylindole (DAPI) to visualize individual cell nuclei and RNA ISH assay was performed according to the Invitrogen ViewRNA ISH Tissue protocol and optimized with the following conditions: heat treatment for ten minutes at 90-95°C and protease digestion at 1:100 dilution for 20 minutes at 40°C. Hybridization was performed for 2h at 40°C using the following species-specific probes: (*Mouse: Cd14* (ThermoFisher, #VB1-3028230), *Cd3e* (ThermoFisher, #VB1-16980), *Cd19* (ThermoFisher, #VB1-3028231), *Rpl13a* (ThermoFisher, #VB1-16196); *Naked mole-rat: Cd14* (ThermoFisher, #VF1-4355532), *Cd3e* (ThermoFisher, #VF1-6000573), *Cd19* (ThermoFisher, #VF1-4359165), *Rpl13a* (ThermoFisher, #VF1-4348271).

### In-situ hybridization data analysis

Whole slide images were generated in fluorescence using Pannoramic SCAN (3D Histech). Image analysis data were generated by automated analysis of whole slide images using ImageDx (Reveal Biosciences), an integrated whole slide image management and automated image analysis workflow. Briefly, each image is first assessed for quality using a focus measurement followed by an accuracy check. All tissue and staining artifacts are digitally excluded from the quantification. The analysis process includes automated identification of tissue, followed by segmentation of regions of interest and then classification of each cell. The total number of cells is determined by detection of DAPI-positive nuclei. Both the number of positive staining cells and the number of positive regions within each cell are recorded. For positive staining cell, the number of cells is provided per each number of puncta, ranging from 1 to 9 and 10 or more (i.e., ten bins).

Using the number of puncta to distinguish between cell types comes with uncertainty since one cannot exclude the possibility that cells with even a single hybridized probe (i.e., puncta = 1) are really negative for the cell-type which this probe is a marker of. Clearly, this uncertainty may affect a test for determining whether mouse and naked mole-rat spleens have different proportions of each of the three cell types targeted by each probe. In order to account for this uncertainty, for each of the ten puncta bins, except for the first, we defined negative cells as those with less than the puncta bin and positive as cells with that number of puncta or higher, and fit a multinomial regression, implemented by the nnet (49) R (43) package, quantifying a species categorical effect, with mouse set as baseline. The uncertainty arising due to selection of a specific puncta bin was propagated by specifying the mean species effect across all puncta bins divided by its standard error as a z-statistic. This allowed obtaining a p-value, which thus served as the statistical significance of the species effect, across all puncta bins, being different from 0.

### RNA isolation

One ml TRIzol reagent (4°C) (Invitrogen, 15596026) was added to ~20mg frozen splenic tissue in a 2ml microcentrifuge tube. A single stainless-steel bead (Qiagen, 69965) was added to each tube and tissue homogenization was performed using a Tissuelyser II (Qiagen) (30Hz for 2mins). RNA was isolated according to manufacturer’s instructions (TRIzol, Invitrogen, 15596026) and resuspended in RNase free water. Contaminating gDNA was removed using the Turbo DNA-free kit (Invitrogen AM1097) according to manufacturer’s instructions. Quantification was performed using the QuBit 4 Fluorometer (Invitrogen) and the Qubit RNA XR assay (Invitrogen, Q33224).

### Whole tissue RNA library preparation, sequencing and data analysis

RNA-seq libraries from the whole splenic tissues were prepared using purified RNA isolated as described above. RNA quality and concentration were assayed using a Fragment Analyzer instrument. RNA-seq libraries were prepared using the TruSeq Stranded Total RNA kit paired with the Ribo-Zero rRNA removal kit (Illumina, San Diego, CA). Libraries were sequenced on an Illumina HiSeq 4000 instrument. Reads were mapped to the naked mole-rat genomic references described above for the scRNA-seq data, using the STAR aligner version 2.5.3a (54), applying the two-round read-mapping approach, with the following parameters: outSAMprimaryFlag = “AllBestScore”, outFilterMultimapNmax = “10”, outFilterMismatchNoverLmax = “0.05”, outFilterIntronMotifs = “RemoveNoncanonical”.

Following read mapping, transcript and gene expression levels were estimated using MMSEQ (55). Transcripts and genes which were minimally distinguishable according to the read data were collapsed using the mmcollapse utility of MMDIFF (56), and the Fragments Per Kilobase Million (FPKM) expression units were converted to Transcripts Per Million (TPM) units. In order to test for differential expression between LPS-challenged and saline tissues, for each species, we used MMDIFF with these two-group design matrices: # M 0, 0, 0, 0; # C 0 0, 0 0, 0 1, 0 1; # P0 1; # P1 0.5 −0.5. Samples were clustered using the heatmap.2 function from the gplots (57) R (43) package, for the purpose of constructing the genes by samples heatmaps for both species. GSEAs were performed for each species, where the inputs to the fgsea function in the fgsea (58) R (43) package were the genes ranked in descending order according to the Bayes Factor and posterior probability reported by MMDIFF, and the gene sets were the Hallmark gene sets of the MSigDB collection (52).

## Author contributions

Conceptualization, HGH, NDR and RB; Methodology, HGH, NDR, NB, KMW, VJ and RB; Investigation, HGH, NDR, PJ, ATI, NB, KMW, VJ, MS, MR, DF, BM, DMI and RB; Acquisition of data, HGH, ATI, DMI; Data curation and analysis, HGH, NDR, NB, KMW, VJ; Writing – original draft, HGH, NDR and DMI; Writing – Review & editing, HGH, NDR, RB; Supervision, MR, VJ and RB.

## Literature Cited

1. Ruby JG, Smith M, Buffenstein R. Naked Mole-Rat mortality rates defy gompertzian laws by not increasing with age. eLife. 2018 24;7.

2. Buffenstein R. The naked mole-rat: a new long-living model for human aging research. J Gerontol A Biol Sci Med Sci. 2005 Nov;60(11):1369–77.

3. Grimes KM, Lindsey ML, Gelfond JAL, Buffenstein R. Getting to the heart of the matter: age-related changes in diastolic heart function in the longest-lived rodent, the naked mole rat. J Gerontol A Biol Sci Med Sci. 2012 Apr;67(4):384–94.

4. O’Connor TP, Lee A, Jarvis JUM, Buffenstein R. Prolonged longevity in naked mole-rats: age-related changes in metabolism, body composition and gastrointestinal function. Comp Biochem Physiol A Mol Integr Physiol. 2002 Nov;133(3):835–42.

5. Labinskyy N, Csiszar A, Orosz Z, Smith K, Rivera A, Buffenstein R, et al. Comparison of endothelial function, O2-* and H2O2 production, and vascular oxidative stress resistance between the longest-living rodent, the naked mole rat, and mice. Am J Physiol Heart Circ Physiol. 2006 Dec;291(6):H2698–2704.

6. Buffenstein R. Negligible senescence in the longest living rodent, the naked mole-rat: insights from a successfully aging species. J Comp Physiol [B]. 2008 May;178(4):439–45.

7. Ferrucci L, Fabbri E. Inflammageing: chronic inflammation in ageing, cardiovascular disease, and frailty. Nat Rev Cardiol. 2018 Sep;15(9):505–22.

8. Nikolich-Žugich J. The twilight of immunity: emerging concepts in aging of the immune system. Nat Immunol. 2018 Jan;19(1):10–9.

9. Chattopadhyay PK, Gierahn TM, Roederer M, Love JC. Single-cell technologies for monitoring immune systems. Nat Immunol. 2014 Feb;15(2):128–35.

10. Zheng GXY, Terry JM, Belgrader P, Ryvkin P, Bent ZW, Wilson R, et al. Massively parallel digital transcriptional profiling of single cells. Nat Commun. 2017 Jan 16;8:14049.

11. Jaitin DA, Kenigsberg E, Keren-Shaul H, Elefant N, Paul F, Zaretsky I, et al. Massively Parallel Single-Cell RNA-Seq for Marker-Free Decomposition of Tissues into Cell Types. Science. 2014 Feb 14;343(6172):776–9.

12. Schaum N, Karkanias J, Neff NF, May AP, Quake SR, Wyss-Coray T, et al. Single-cell transcriptomics of 20 mouse organs creates a Tabula Muris. Nature. 2018 Oct 1;562(7727):367–72.

13. Dutta P, Hoyer FF, Grigoryeva LS, Sager HB, Leuschner F, Courties G, et al. Macrophages retain hematopoietic stem cells in the spleen via VCAM-1. J Exp Med. 2015 Apr 6;212(4):497–512.

14. Sehgal A, Price J, Man B, Young M. Loss of circadian behavioral rhythms and per RNA oscillations in the Drosophila mutant timeless. Science. 1994 Mar 18;263(5153):1603–6.

15. Leman AR, Dheekollu J, Deng Z, Lee SW, Das MM, Lieberman PM, et al. Timeless preserves telomere length by promoting efficient DNA replication through human telomeres. Cell Cycle Georget Tex. 2012 Jun 15;11(12):2337–47.

16. Cao A, Moi P. Regulation of the globin genes. Pediatr Res. 2002 Apr;51(4):415–21.

17. Sankaran VG, Xu J, Orkin SH. Advances in the understanding of haemoglobin switching. Br J Haematol. 2010 Apr;149(2):181–94.

18. Takakuwa Y. Regulation of red cell membrane protein interactions: implications for red cell function. Curr Opin Hematol. 2001 Mar;8(2):80–4.

19. Palis J, Malik J, McGrath KE, Kingsley PD. Primitive erythropoiesis in the mammalian embryo. Int J Dev Biol. 2010;54(6–7):1011–8.

20. Crinier A, Milpied P, Escalière B, Piperoglou C, Galluso J, Balsamo A, et al. High-Dimensional Single-Cell Analysis Identifies Organ-Specific Signatures and Conserved NK Cell Subsets in Humans and Mice. Immunity. 2018 Nov 1;

21. Dwyer DF, Barrett NA, Austen KF, Immunological Genome Project Consortium. Expression profiling of constitutive mast cells reveals a unique identity within the immune system. Nat Immunol. 2016 Jul;17(7):878–87.

22. Guethlein LA, Norman PJ, Hilton HG, Parham P. Co-evolution of MHC class I and variable NK cell receptors in placental mammals. Immunol Rev. 2015;267(1):259–82.

23. Kelley J, Walter L, Trowsdale J. Comparative genomics of natural killer cell receptor gene clusters. PLoS Genet. 2005 Aug;1(2):129–39.

24. Keane M, Craig T, Alföldi J, Berlin AM, Johnson J, Seluanov A, et al. The Naked Mole Rat Genome Resource: facilitating analyses of cancer and longevity-related adaptations. Bioinforma Oxf Engl. 2014 Dec 15;30(24):3558–60.

25. Parham P, Moffett A. Variable NK cell receptors and their MHC class I ligands in immunity, reproduction and human evolution. Nat Rev Immunol. 2013 Feb;13(2):133–44.

26. Schenkel AR, Kingry LC, Slayden RA. The ly49 gene family. A brief guide to the nomenclature, genetics, and role in intracellular infection. Front Immunol. 2013;4:90.

27. Debout C, Quillec M, Izard J. Natural killer activity of Kurloff cells: a direct demonstration on purified Kurloff cell suspensions. Cell Immunol. 1984 Sep;87(2):674–7.

28. Jara LF, Sánchez JM, Alvarado H, Nassar-Montoya F. Kurloff cells in peripheral blood and organs of wild capybaras. J Wildl Dis. 2005 Apr;41(2):431–4.

29. Wright SD, Ramos RA, Tobias PS, Ulevitch RJ, Mathison JC. CD14, a receptor for complexes of lipopolysaccharide (LPS) and LPS binding protein. Science. 1990 Sep 21;249(4975):1431–3.

30. Schumann RR, Leong SR, Flaggs GW, Gray PW, Wright SD, Mathison JC, et al. Structure and function of lipopolysaccharide binding protein. Science. 1990 Sep 21;249(4975):1429–31.

31. Shalek AK, Satija R, Shuga J, Trombetta JJ, Gennert D, Lu D, et al. Single-cell RNA-seq reveals dynamic paracrine control of cellular variation. Nature. 2014 Jun 19;510(7505):363–9.

32. La Manno G, Soldatov R, Zeisel A, Braun E, Hochgerner H, Petukhov V, et al. RNA velocity of single cells. Nature. 2018;560(7719):494–8.

33. Vivier E, Tomasello E, Baratin M, Walzer T, Ugolini S. Functions of natural killer cells. Nat Immunol. 2008 May;9(5):503–10.

34. Long EO, Kim HS, Liu D, Peterson ME, Rajagopalan S. Controlling natural killer cell responses: integration of signals for activation and inhibition. Annu Rev Immunol. 2013;31:227–58.

35. Pavlovich SS, Lovett SP, Koroleva G, Guito JC, Arnold CE, Nagle ER, et al. The Egyptian Rousette Genome Reveals Unexpected Features of Bat Antiviral Immunity. Cell. 2018 May 17;173(5):1098–1110.e18.

36. Hammond JA, Guethlein LA, Abi-Rached L, Moesta AK, Parham P. Evolution and survival of marine carnivores did not require a diversity of killer cell Ig-like receptors or Ly49 NK cell receptors. J Immunol Baltim Md 1950. 2009 Mar 15;182(6):3618–27.

37. Cho H, Soundrarajan N, Le Van Chanh Q, Jeon H, Cha S-Y, Kang M, et al. The novel cathelicidin of naked mole rats, Hg-CATH, showed potent antimicrobial activity and low cytotoxicity. Gene. 2018 Nov;676:164–70.

38. Ross-Gillespie A, O’Riain MJ, Keller LF. VIRAL EPIZOOTIC REVEALS INBREEDING DEPRESSION IN A HABITUALLY INBREEDING MAMMAL. Evolution. 2007 Sep;61(9):2268–73.

39. Artwohl J, Ball-Kell S, Valyi-Nagy T, Wilson SP, Lu Y, Park TJ. Extreme susceptibility of African naked mole rats (Heterocephalus glaber) to experimental infection with herpes simplex virus type 1. Comp Med. 2009 Feb;59(1):83–90.

40. Mudge JM, Harrow J. Creating reference gene annotation for the mouse C57BL6/J genome assembly. Mamm Genome Off J Int Mamm Genome Soc. 2015 Oct;26(9–10):366–78.

41. Butler A, Hoffman P, Smibert P, Papalexi E, Satija R. Integrating single-cell transcriptomic data across different conditions, technologies, and species. Nat Biotechnol. 2018 Jun;36(5):411–20.

42. Erichson NB, Voronin S, Brunton SL, Kutz JN. Randomized Matrix Decompositions using R. ArXiv160802148 Cs Stat [Internet]. 2016 Aug 6 [cited 2019 Feb 26]; Available from: http://arxiv.org/abs/1608.02148

43. R Core Team. R: A language and environment for statistical computing. 2014.

44. Waltman L, van Eck NJ. A smart local moving algorithm for large-scale modularity-based community detection. Eur Phys J B [Internet]. 2013 Nov [cited 2018 Dec 4];86(11). Available from: http://link.springer.com/10.1140/epjb/e2013-40829-0

45. Benjamini Y, Drai D, Elmer G, Kafkafi N, Golani I. Controlling the false discovery rate in behavior genetics research. Behav Brain Res. 2001 Nov 1;125(1–2):279–84.

46. Altschul SF, Madden TL, Schäffer AA, Zhang J, Zhang Z, Miller W, et al. Gapped BLAST and PSI-BLAST: a new generation of protein database search programs. Nucleic Acids Res. 1997 Sep 1;25(17):3389–402.

47. Emms DM, Kelly S. OrthoFinder: solving fundamental biases in whole genome comparisons dramatically improves orthogroup inference accuracy. Genome Biol. 2015 Aug 6;16:157.

48. Frankish A, Diekhans M, Ferreira A-M, Johnson R, Jungreis I, Loveland J, et al. GENCODE reference annotation for the human and mouse genomes. Nucleic Acids Res. 2019 Jan 8;47(D1):D766–73.

49. Venables WN, Ripley BD, Venables WN. Modern applied statistics with S. 4th ed. New York: Springer; 2002. 495 p. (Statistics and computing).

50. Bates D, Mächler M, Bolker B, Walker S. Fitting Linear Mixed-Effects Models Using lme4. J Stat Softw [Internet]. 2015[cited 2019 Feb 26];67(1). Available from: http://www.jstatsoft.org/v67/i01/

51. The Gene Ontology Consortium. The Gene Ontology Resource: 20 years and still GOing strong. Nucleic Acids Res. 2019 Jan 8;47(D1):D330–8.

52. Liberzon A, Subramanian A, Pinchback R, Thorvaldsdottir H, Tamayo P, Mesirov JP. Molecular signatures database (MSigDB) 3.0. Bioinformatics. 2011 Jun 15;27(12):1739–40.

53. Ashburner M, Ball CA, Blake JA, Botstein D, Butler H, Cherry JM, et al. Gene ontology: tool for the unification of biology. The Gene Ontology Consortium. Nat Genet. 2000 May;25(1):25–9.

54. Dobin A, Davis CA, Schlesinger F, Drenkow J, Zaleski C, Jha S, et al. STAR: ultrafast universal RNA-seq aligner. Bioinforma Oxf Engl. 2013 Jan 1;29(1):15–21.

55. Turro E, Su S-Y, Gonçalves Â, Coin LJM, Richardson S, Lewin A. Haplotype and isoform specific expression estimation using multi-mapping RNA-seq reads. Genome Biol. 2011;12(2):R13.

56. Turro E, Astle WJ, Tavaré S. Flexible analysis of RNA-seq data using mixed effects models. Bioinforma Oxf Engl. 2014 Jan 15;30(2):180–8.

57. Warnes GR, Bolker B, Bonebakker L, Gentleman R, Huber W, Liaw A, et al. gplots: Various R programming tools for plotting data. R Package Version. 2009;2(4):1.

58. Sergushichev A. An algorithm for fast preranked gene set enrichment analysis using cumulative statistic calculation. bioRxiv [Internet]. 2016 Jun 20 [cited 2019 Jan 2]; Available from: http://biorxiv.org/lookup/doi/10.1101/060012

