## Supplementary Table and Figure Legends for "Single-cell transcriptomics of the naked mole-rat reveals unexpected features of mammalian immunity"

### Supplementary Figure Legends

**Supplementary Figure 1:** *Uniform representation across samples in cluster composition in the spleen single-cell RNA-sequencing data*

**a -** Stacked bar chart showing the proportion (%) of mouse spleen cells from four animals that corresponds to a given cluster/cell type from the first clustering iteration. Each pair of color shades represents cells from the duplicate samples from each of the four mice. **b -** Stacked bar chart showing the proportion (%) of mouse spleen cells from four animals that corresponds to a given cluster/cell type at convergence of the clustering. Each pair of color shades represents cells from the duplicate samples from each of the four mice. **c -** Stacked bar chart showing the proportion (%) of naked mole-rat spleen cells from four animals that corresponds to a given cluster/cell type from the first clustering iteration. Each pair of color shades represents cells from the duplicate samples from each of four naked mole-rats. **d -** Stacked bar chart showing the proportion (%) of naked mole-rat spleen cells from four animals that corresponds to a given cluster/cell type at convergence of the clustering. Each pair of color shades represents cells from the duplicate samples from each of four naked mole-rats.

**Supplementary Figure 2:** *A transcriptional map of immune cells from the mouse and naked mole-rat spleen at single-cell resolution from the converged clustering*

**a -** Mouse spleen cell types clustered to convergence and visualized by uniform manifold approximation and projection (UMAP). Colors indicate clusters that correspond to the listed cell types. **b** – Naked mole-rat spleen cell types clustered to convergence and visualized by uniform manifold approximation and projection (UMAP). Colors indicate clusters that correspond to the listed cell types. **c -** Heatmap showing the gene expression in mouse spleen cells in each of twenty-six clusters that correspond to the listed cell types. Selected marker genes are listed to the left. Clusters for which an assignment was not possible are listed as unassigned. **d -** Heatmap showing the gene expression in naked mole-rat spleen cells in each of twenty-two clusters that correspond to the listed cell types. Selected marker genes are listed to the left. Clusters for which an assignment was not possible are listed as unassigned.

**Supplementary Figure 3:** *Naked mole-rat spleen erythroid cell sub-clusters reveal the presence of embryonic hemoglobin*

Naked mole-rat cell clusters visualized by UMAP showing the relative expression of the $\beta$ (**a**), $\alpha$ (**b**), $\varepsilon$ (**c**), $\zeta$ (**d**), $\theta$ (**e**) and $\gamma$ (**f**) globin genes and of the *glycophorin c* gene (*Gypc*) (**g**), the gene encoding the erythrocyte membrane protein band 4.1 (*Epb41*) (**h**), *Epb41* versus *Gypc* relative expression (scaled UMIs) among sub-cluster 2 cells (**i**), *Epb41* versus *Gypc* relative expression among sub-cluster 9 cells (**j**), and of the erythropoietin receptor gene (*Epor*) (**k**), *Bcl-xL* (*Bcl2l1*) (**l**), *Bcl2l1* versus *Epor* relative expression among sub-cluster 2 cells (**m**) and *Bcl2l1* versus *Epor* relative expression among sub-cluster 9 cells (**n**). In the gene-gene scatter plots, r denotes the Pearson correlation coefficient and p the associated p-value of the coefficient being different from zero. **o -** Violin plots showing the distributions of the numbers of genes sequenced in myeloid lineage cell subsets in the naked mole-rat spleen. To the right is shown the estimated cell-type effect sizes (relative to sub-cluster 10) (asterisks mark adjusted p-value < 0.05).

**Supplementary Figure 4:** *A lack of annotated natural killer cell marker genes is not likely to account for the absence of an identifiable natural killer cell cluster in the naked mole-rat spleen*

**a** – Naked mole-rat cell clusters visualized by UMAP showing the relative expression of *Nkg7*, *Gzma*, *Ccl5*, *Tpsab1*, *Cma1*, and *Fcer1a.* **b** – Mouse spleen cell clusters obtained using a complete gene list and visualized by UMAP. Colors indicate clusters that correspond to the listed cell types. Shown below is a table that lists the number and fraction (%) of cells in each cluster. **c** – Mouse spleen cell clusters obtained using a gene list depleted of NK cell markers and visualized by UMAP. Colors indicate clusters that correspond to the listed cell types. Shown below is a table that lists the number and fraction (%) of cells in each cluster.

**Supplementary Figure 5:** *A transcriptionally defined cluster of cells corresponding to natural killer cells is not found in circulating immune cells from the naked mole-rat*

**a** – Schematic showing the workflow in which single-cell suspensions are derived from circulating immune cells of four (two male [M], two female [F]) C57BL/6 mice (upper panel, grey) or four naked mole-rats (two male [M], two female [F]) (lower panel, orange). **b** – Bar chart showing the number of cells sequenced from each of four mice (upper panel, grey) and three naked mole-rats (lower panel, orange) [One naked mole-rat sample was not sequenced due to a microfluidic failure during the emulsion generation]. **c** – Violin plots showing the number of genes sequenced per cell from each of four mice (upper panel, grey) and three naked mole-rats (lower panel, orange). **d** – Violin plots showing the number of unique molecular identifiers (UMIs) sequenced per cell from each of four mice (upper panel, grey) and three naked mole-rats (lower panel, orange). **e** – Mouse circulating immune cell clusters visualized by UMAP. Colors indicate clusters that correspond to the listed cell types. The relative proportions of each cell type are shown in a stacked bar chart to the right. **f** – Naked mole-rat spleen cell clusters visualized by UMAP. Colors indicate clusters that correspond to the listed cell types. The relative proportions of each cell type are shown in a stacked bar chart to the left. **g** – Heatmap showing the gene expression in mouse circulating immune cells in each of six clusters that correspond to the listed cell type. Selected marker genes are listed to the left. **h** – Heatmap showing the gene expression in naked mole-rat circulating immune cells in each of nine clusters that correspond to the listed cell type. Selected marker genes are listed to the left.

**Supplementary Figure 6:** *A lack of annotated natural killer cell marker genes is not likely to account for the absence of an identifiable natural killer cell cluster in circulating immune cells from the naked mole-rat*

**a** – Mouse spleen cell clusters obtained using a gene list depleted of NK cell markers and visualized by UMAP. Colors indicate clusters that correspond to the listed cell types. **b -** Mouse circulating immune cell converged clusters obtained using a gene list depleted of NK cell markers and visualized by UMAP. Colors indicate clusters that correspond to the listed cell types.

**Supplementary Figure 7:** *Sequence analysis of immunoglobulin-like and lectin-like receptors encoded in the putative naked mole-rat natural killer cell receptor complexes*

Amino acid sequences encoded by the *Klra* (**a**), *Cd94* (**b**) and *Nkg2* (**c**) genes encoded in the putative naked mole-rat *NKC*. Shown for each protein sequence is the region predicted to form the extracellular region (green), the intracellular region (blue) and the transmembrane region (red). Annotations below each entry indicate features that are absent from that sequence. Immunoreceptor tyrosine inhibitory motifs (ITIMs) and double cysteine residues are shown in pink.

**Supplementary Figure 8:** *The naked mole-rat has only one expressed major histocompatibility complex class I receptor gene*

**a** – Naked mole-rat spleen cell clusters visualized by UMAP showing the relative expression of *Klra1 (Ly49* family) and *Klrd1* (*Cd94*). **b** – Mouse cell clusters visualized by UMAP showing the relative expression of *Klra1 (Ly49* family*), Klrd1* (*Cd94*) and *Klrc1* (*Nkg2a*).

**Supplementary Figure 9:** *The naked mole-rat spleen does not contain Foa-Kurloff cells*

**a, b -** Representative images of Periodic acid-Schiff (PAS) stained sections of naked mole-rat spleen. Scale bar = 500µm.

**Supplementary Figure 10:** *Whole-spleen RNA sequencing shows broadly conserved affected biological pathways between mice and naked mole-rats following lipopolysaccharide challenge*

**a** – Genes × samples scaled ln(TPM) heatmap in spleens from control (saline, n=2) and lipopolysaccharide (LPS, n=2) challenged mice. Selected LPS upregulated genes are shown to the left. **b** – Genes × samples scaled ln(TPM) heatmap in spleens from control (saline, n=2) and lipopolysaccharide (LPS, n=2) challenged naked mole-rats. Selected LPS upregulated genes are shown to the right. **c** – Volcano plot showing the posterior probability of the estimated effect size (ln(LPS/saline) ln(TPM) fold-change) being different from zero (y-axis) versus the estimated effect size (x-axis) in the mouse data (saline n=2; LPS n=2). Each point is a gene and the color code follows the posterior probability gradient. **d -** Volcano plot showing the posterior probability of the estimated effect size (ln(LPS/saline) ln(TPM) fold-change) being different from zero (y-axis) versus the estimated effect size (x-axis) in the naked mole-rat data (saline n=2; LPS n=2). Each point is a gene and the color code follows the posterior probability gradient. **e** – Bar chart showing the hallmark gene sets enriched in genes with strong expression changes following LPS challenge in mice. The x-axis reports the $\log_{10}$ adjusted p-value (q-value) of the GSEA, signed by the direction of the effect (up- and down -regulation in LPS relative to saline: positive and yellow and negative and cyan, respectively). Vertical dashed lines represent adjusted p-value = 0.1. **f** – Bar chart showing the hallmark gene sets enriched in genes with strong expression changes following LPS challenge in naked mole-rats. The x-axis reports the $\log_{10}$ adjusted p-value (q-value) of the GSEA, signed by the direction of the effect (up- and down -regulation in LPS relative to saline: positive and yellow and negative and cyan, respectively). Vertical dashed lines represent adjusted p-value = 0.1.

**Supplementary Figure 11:** *A transcriptional map of circulating immune cells from the mouse and naked mole-rat at single-cell resolution from control and lipopolysaccharide treated animals*
**a -** Schematic showing the experimental outline in which four C57BL/6 mice are assigned to either saline control (n=2) or LPS-treated (n=2) groups. After four hours, single-cell suspensions were derived from circulating immune cells of each animal and subjected to single-cell RNA sequencing. **b -** Schematic showing the experimental outline in which four naked mole-rats are assigned to either saline control (n=2) or LPS-treated (n=2) groups. After four hours, single-cell suspensions were derived from circulating immune cells of each animal and subjected to single-cell RNA sequencing. **c -** Mouse circulating-immune-cell clusters from saline control (left panel) and LPS-treated (right panel) visualized by UMAP. Colors indicate clusters that correspond to the listed cell types. **d -** Naked mole-rat circulating-immune-cell clusters from saline control (left panel) and LPS-treated (right panel) visualized by UMAP. Colors indicate clusters that correspond to the listed cell types. **e -** Stacked bar charts showing the relative proportions (%) of the listed cell types in saline control and LPS-treated mice (upper panel) with the ln(LPS/saline) cell-count ratios are shown as effect sizes (lower panel). **f -** Stacked bar charts showing the relative proportions (%) of the listed cell types in saline control and LPS-treated naked mole-rats (upper panel) with the ln(LPS/saline) cell-count ratios are shown as effect sizes (lower panel). **g -** Heatmap showing the gene expression in mouse WBC from saline control (-) and LPS-treated (+) animals in each of six clusters (mean across the cells of each cluster) that correspond to the listed cell type. Selected marker genes are listed to the left. **h -** Heatmap showing the gene expression in naked mole-rat circulating immune cells from saline control (-) and LPS-treated (+) animals in each of six clusters (mean across the cells of each cluster) that correspond to the listed cell types. Selected marker genes are listed to the left.

**Supplementary Figure 12:** *Cell specific responses to lipopolysaccharide challenge in circulating immune cells from the naked mole-rat and mouse*
**a -** Selected density plots showing the expression distribution of *Nfkbia* from saline control (orange) and LPS-challenged (red) mice across the cells of five representative cell types. Insets

show the estimated treatment expression and expression-induction effect sizes (asterisks mark adjusted p-value < 0.05). **b** - Selected density plots showing the expression distribution of *Nfkbia* from saline control (orange) and LPS-challenged (red) naked mole-rats across the cells of five representative cell types. Insets show the estimated treatment expression and expression-induction effect sizes (LPS relative to saline) (asterisks mark adjusted p-value < 0.05). **c** - Heatmap showing the estimated treatment-effect statistics for expression change. Selected marker genes are shown to the left. **d** - Heatmap showing the estimated treatment-effect statistics for expression-induction change. Selected marker genes are shown to the right. **e** - Bar chart showing the gene set enrichment analyses (GSEAs) of the intra-species treatment effect on expression change in mouse NK cells. The x-axis reports the $\log_{10}$ adjusted p-value (q-value) of the GSEA. The $\log_{10}$ adjusted p-value is signed and color-coded by the direction of the effect (up- and down -regulation in LPS relative to saline: yellow and cyan, respectively). Black vertical dashed lines correspond to an adjusted p-value of 0.1. **f -** Heatmap showing GSEAs of the intra-species treatment effects on expression-induction change in mice across the listed cell types. The $\log_{10}$ adjusted p-value (q-value) of the GSEA is indicated by the shade of the color and the colors encode the direction of the effect (up- and down -regulation in LPS relative to saline: yellow and cyan, respectively). Gene sets with an adjusted p-value of < 0.1 are indicated with an asterisk. **g -** Heatmap showing GSEAs of the intra-species treatment effect on expression-induction change in naked mole-rats across the listed cell types. The $\log_{10}$ adjusted p-value (q-value) of the GSEA is indicated by the shade of the color and the colors encode the direction of the effect (up- and down -regulation in LPS relative to saline: yellow and cyan, respectively). Gene sets with an adjusted p-value of < 0.1 are indicated with an asterisk. **h -** Clusters representing *Ltf*-high neutrophils (pink) and neutrophils (yellow) from saline control animals visualized by UMAP. Overlaid on each cluster are arrows that represent the ratio of spliced to unspliced RNA (RNA velocity). **i -** Clusters representing *Ltf*-high neutrophils (pink) and neutrophils (yellow) from LPS-challenged animals visualized by UMAP. Overlaid on each cluster are arrows that represent the ratio of spliced to unspliced RNA (RNA velocity). **j -** Bar chart showing GSEAs comparing *Ltf*-high neutrophils to neutrophils from LPS-challenged naked mole-rats. The x-axis reports the $\log_{10}$ adjusted p-value (q-value) of the GSEA. The $\log_{10}$ adjusted p-value is signed and color-coded by the direction of the effect (up- and down -regulation in *Ltf*-high neutrophils relative to neutrophils: yellow and cyan, respectively). Black vertical dashed lines correspond to an adjusted p-value of 0.1.

**Supplementary Table Legends**

**Supplementary Table 1:** Table showing the number of unique molecular identifiers (UMIs) and genes recorded per spleen cell captured by single cell RNA sequencing in four C57BL/6 mice. Cells from each mouse spleen were sequenced in duplicate and designated *mouse_1.1, 1.2; 2.1, 2.2; 3.1, 3.2; 4.1, 4.2.* For each cell is also listed its parental and converged cluster designation (see *Methods*).

**Supplementary Table 2:** Table showing the number of unique molecular identifiers (UMIs) and genes recorded per spleen cell captured by single cell RNA sequencing in four naked mole-rats. Cells from each naked mole-rat spleen were sequenced in duplicate and designated *nmr_1.1, 1.2; 2.1, 2.2; 3.1, 3.2; 4.1, 4.2.* For each cell is also listed its first-iteration and converged cluster (see *Methods*) designation and assigned cell type.

**Supplementary Table 3:** Table showing the identity of marker genes used to cluster spleen cells from four C57BL/6 mice. For each gene is listed its Ensembl gene ID, symbol, description, biotype, the identity of the converged cluster in which it is highly expressed, and the cell type assigned to that cluster.

**Supplementary Table 4:** Table showing the identity of marker genes used to cluster spleen cells from four naked mole-rats. For each gene is listed its NCBI gene ID, symbol, the identity of the converged cluster in which it is highly expressed, and the cell type assigned to that cluster.

**Supplementary Table 5:** Table showing the results of quantification of in-situ hybridization studies from the spleen of four C57BL/6 mice (designated Mouse_5-8) and four naked mole-rats (designated NMR_5-8). For each animal is shown the results of staining with probes targeting *Cd19, Cd3e* and *Cd14.* For each spleen/probe combination is shown the total number of cells quantified, as well as the number of positively staining puncta detected per cell (range 1-10+) and the total area quantified.

**Supplementary Table 6:** Table showing the results comparing the numbers of genes across myeloid lineage cells of the naked mole-rat spleen with cells of erythroid sub-cluster 10). For each myeloid lineage cell type contrasted with cells of erythroid sub-cluster 10, shown are the estimated effect size, error of effect size, p-value and adjusted p-value of the effect size.

**Supplementary Table 7:** Table showing the number and fraction of mouse spleen cells from the first-iteration clusters (1 – 6), obtained by re-clustering in the absence of NK cell marker genes, that are correctly re-assigned into that same cluster obtained from the fully annotated data.

**Supplementary Table 8:** Table showing the number of unique molecular identifiers (UMIs) and genes recorded per circulating immune cell captured by single cell RNA sequencing in four C57BL/6 mice. Samples are designated *mouse.CIC_1-4*. For each cell is also listed its first-iteration and converged cluster (see *Methods*) designation and assigned cell type.

**Supplementary Table 9:** Table showing the number of unique molecular identifiers (UMIs) and genes recorded per circulating immune cell captured by single cell RNA sequencing in three naked mole-rats. Samples are designated *nmr.CIC_2-4* (sample 1 filtered due to microfluidic failure). For each cell is also listed its first-iteration and converged cluster (see *Methods*) designation and assigned cell type.

**Supplementary Table 10:** Table showing the number and fraction of mouse circulating immune cells from the first-iteration clusters (1 – 5), obtained by re-clustering in the absence of NK cell marker genes, that are correctly re-assigned into that same cluster obtained from the fully annotated data.

**Supplementary Table 11:** Table showing the number and fraction of mouse circulating immune cells from the converged clusters (1 – 9), obtained by re-clustering in the absence of NK cell marker genes, that are correctly re-assigned into that same cluster obtained from the fully annotated data.

**Supplementary Table 12:** Table showing the genes in the human, mouse, rat, guinea pig and naked mole-rat leukocyte receptor and natural killer cell receptor complexes (*NKC* and *LRC* respectively). For each species, given are its Ensembl gene ID (NCBI for the naked mole-rat), its gene name, scaffold name, scaffold start and end coordinates and predicted biotype. In addition, for each gene its complex, gene family as well as its orientation relative to the human genome are indicated.

**Supplementary Table 13:** Table showing the differential expression of genes in whole spleen tissue between saline control (n=2) and LPS challenged (n=2) mice.

**Supplementary Table 14***:* Table showing the differential expression of genes in whole spleen tissue between saline control (n=2) and LPS challenged (n=2) naked mole-rats.

**Supplementary Table 15** Table showing the results of hallmark gene set enrichment analyses of genes differentially expressed in whole spleen tissue between saline control (n=2) and LPS challenged (n=2) mice.

**Supplementary Table 16***:* Table showing the results of hallmark gene set enrichment analyses of genes differentially expressed in whole spleen tissue between saline control (n=2) and LPS challenged (n=2) naked mole-rats.

**Supplementary Table 17**: Table showing the results of hallmark gene set enrichment analyses of genes differentially expressed in splenic NK cells between saline control (n=2) and LPS challenged (n=2) mice.

**Supplementary Table 18:** Table showing the results of hallmark gene set enrichment analyses of genes differentially induced in splenic monocytes, neutrophils, macrophages, B- and T-cells between saline control (n=2) and LPS challenged (n=2) mice.

**Supplementary Table 19:** Table showing the results of hallmark gene set enrichment analyses of genes differentially induced in splenic monocytes and mast cells between saline control (n=2) and LPS challenged (n=2) naked mole-rats.

**Supplementary Table 20:** Table showing the results of Gene-Ontology (GO) Biological pathway enrichment gene set enrichment analyses of genes differentially expressed between splenic neutrophils and *Ltf*-high neutrophils in saline control (n=2) naked mole-rats.

**Supplementary Table 21:** Table showing the results of Gene-Ontology (GO) Biological pathway enrichment gene set enrichment analyses of genes differentially expressed between splenic neutrophils and *Ltf*-high neutrophils in LPS challenged (n=2) naked mole-rats.

**Supplementary Table 22:** Table showing the results of hallmark gene set enrichment analyses of genes differentially expressed in circulating NK cells between saline control (n=2) and LPS challenged (n=2) mice.

**Supplementary Table 23:** Table showing the results of Hallmark gene set enrichment analyses of genes differentially induced in circulating monocytes, neutrophils, B- and T-cells between saline control (n=2) and LPS challenged (n=2) mice.

**Supplementary Table 24:** Table showing the results of hallmark gene set enrichment analyses of genes differentially induced in circulating B- and T-cells between saline control (n=2) and LPS challenged (n=2) naked mole-rats.

**Supplementary Table 25:** Table showing the results of Gene-Ontology (GO) Biological pathway gene set enrichment analyses of genes differentially expressed between circulating neutrophils and *Ltf*-high neutrophils in LPS challenged (n=2) naked mole-rats.

**Supplementary Table 26:** Table showing the summary statistics of single-cell RNA sequencing data from the circulating immune cells and spleen of mice and naked mole-rats in this study.

**Supplementary Table 27:** Table showing the one-to-one gene orthologs between mouse and naked mole-rat.

**Supplementary Table 28:** Table showing the matched cell types in mouse and naked mole-rat (separately for each species) spleen control saline and LPS-challenge conditions.

**Supplementary Table 29:** Table showing the matched cell types in mouse and naked mole-rat (separately for each species) circulating immune cells.

**Supplementary Table 30:** Table showing the matched cell types between mouse and naked mole-rat spleen control saline and LPS-challenge conditions used for constructing Figures 5c-d.

**Supplementary Table 31:** Table showing the matched cell types between mouse and naked mole-rat circulating immune cells control saline and LPS-challenge conditions used for constructing Figures S12c-d.
