## Supplementary figures and images for "Single-cell transcriptomics of the naked mole-rat reveals unexpected features of mammalian immunity"

### FigureS1

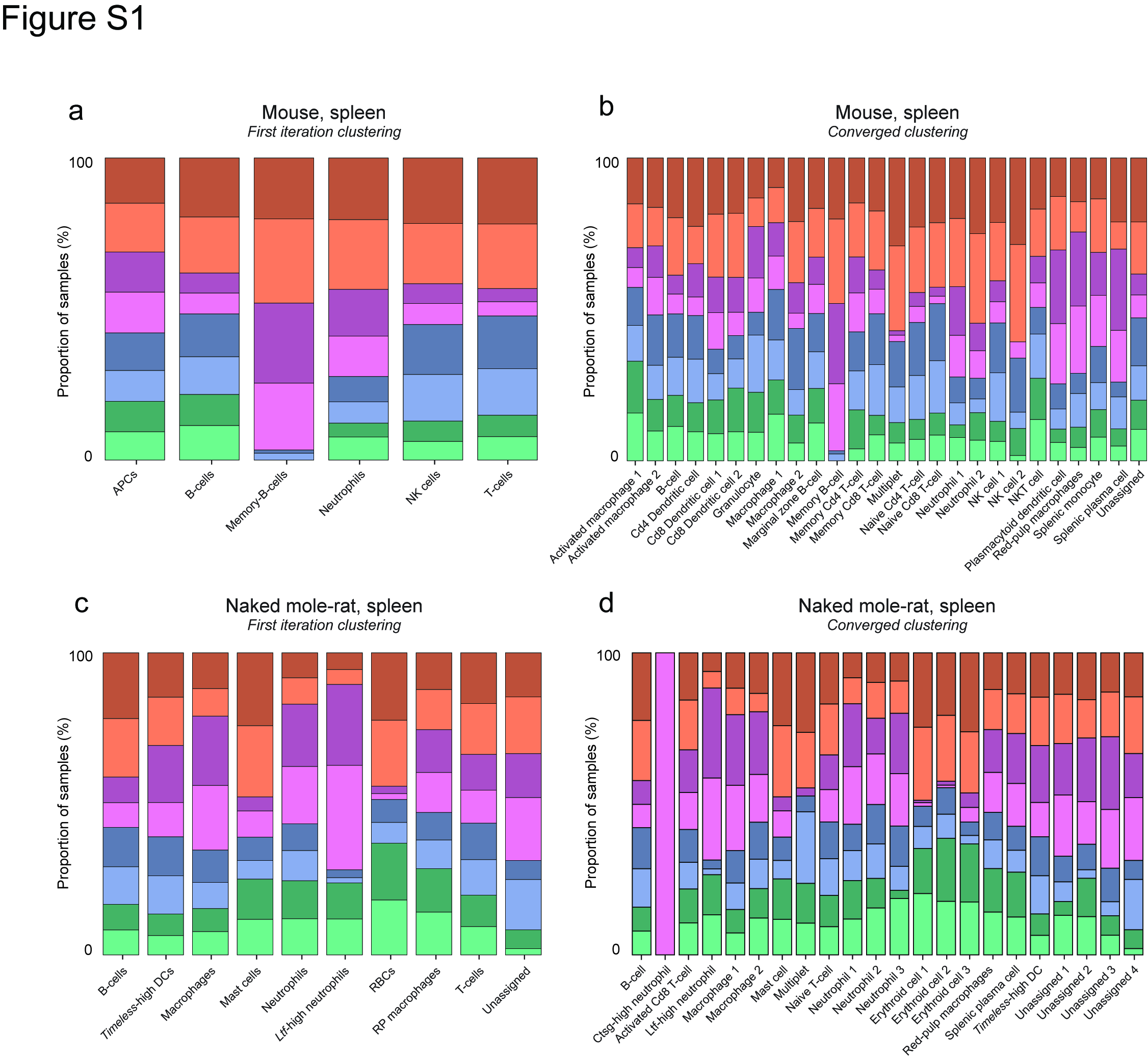

### FigureS2

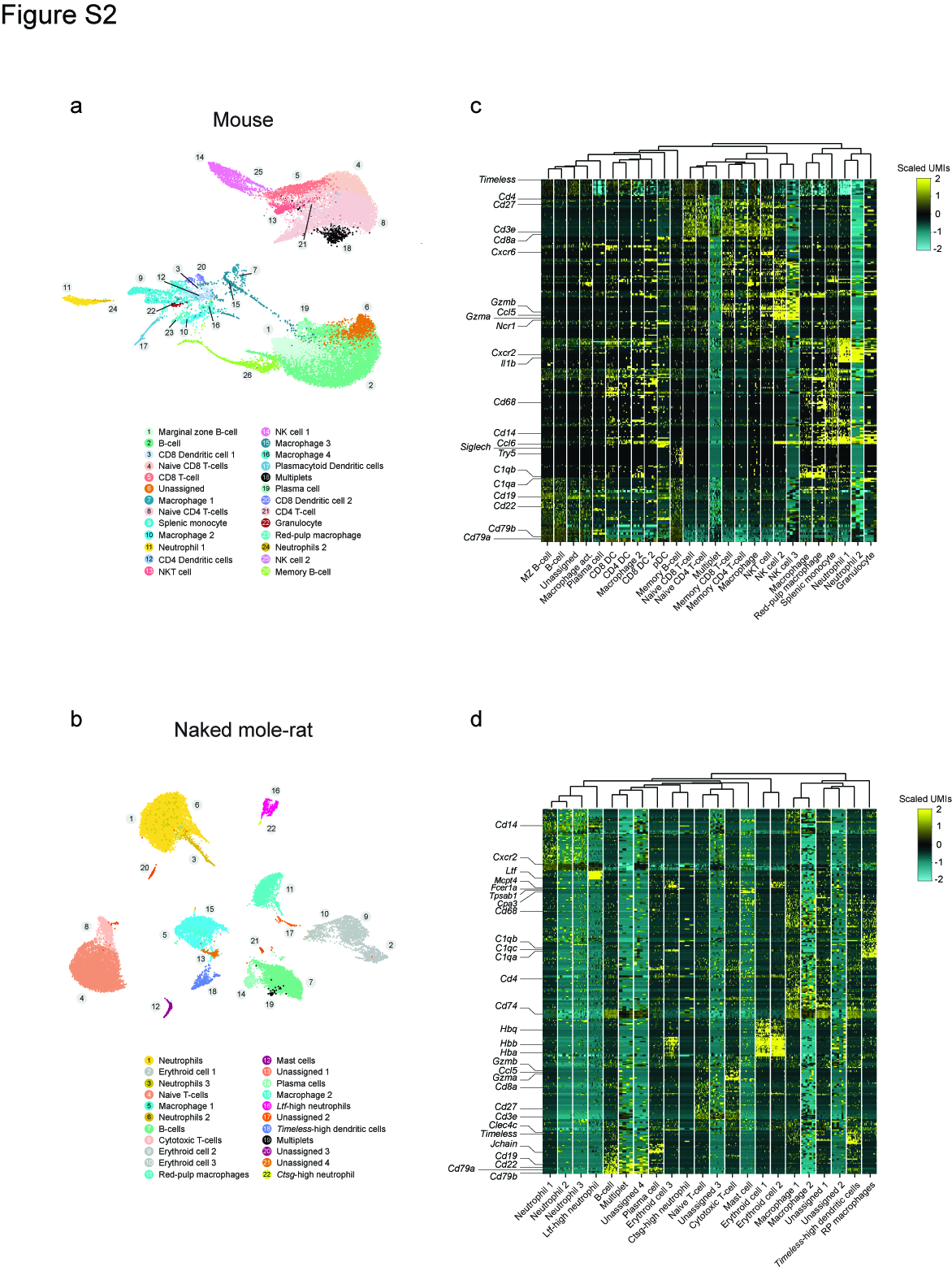

### FigureS3

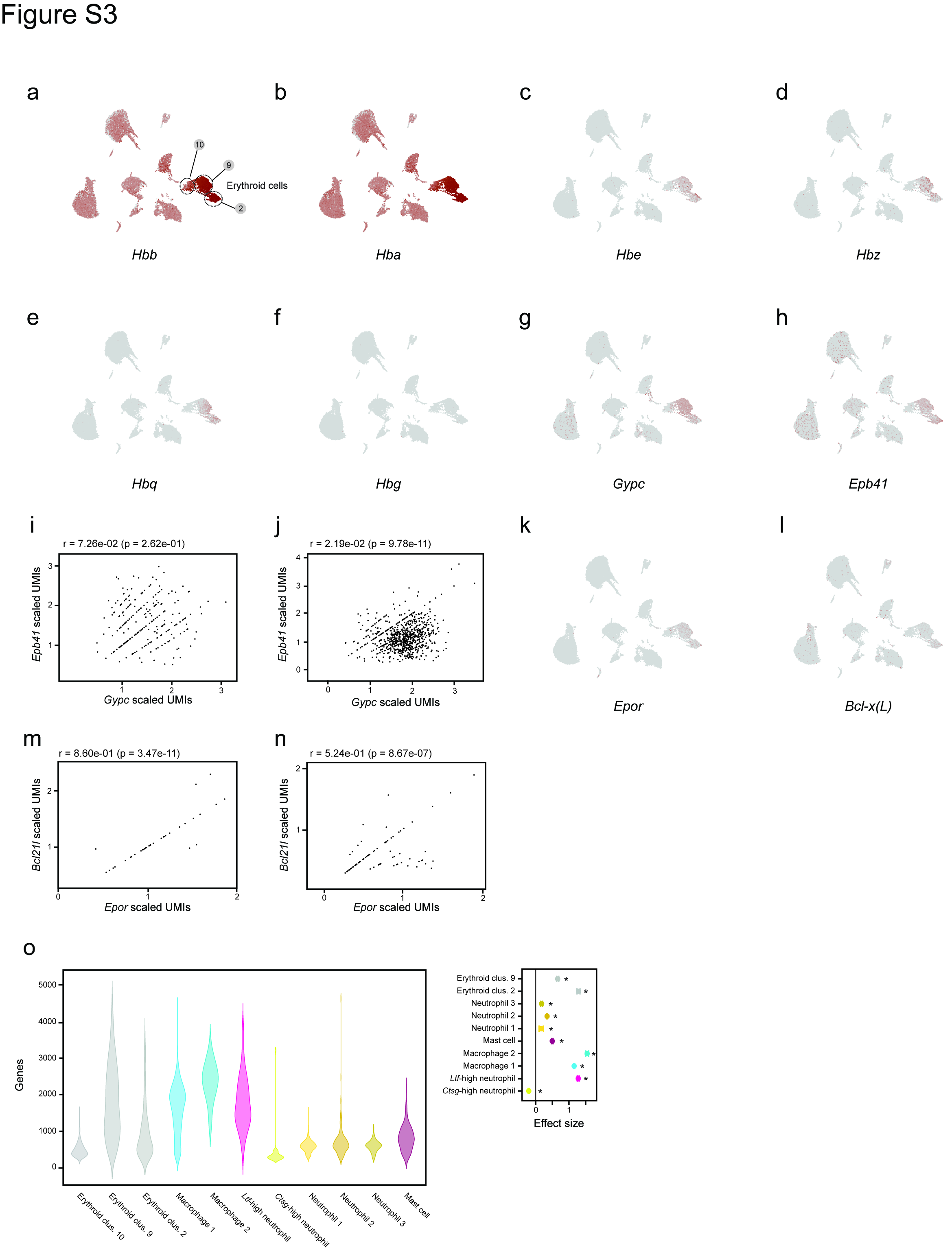

### FigureS4

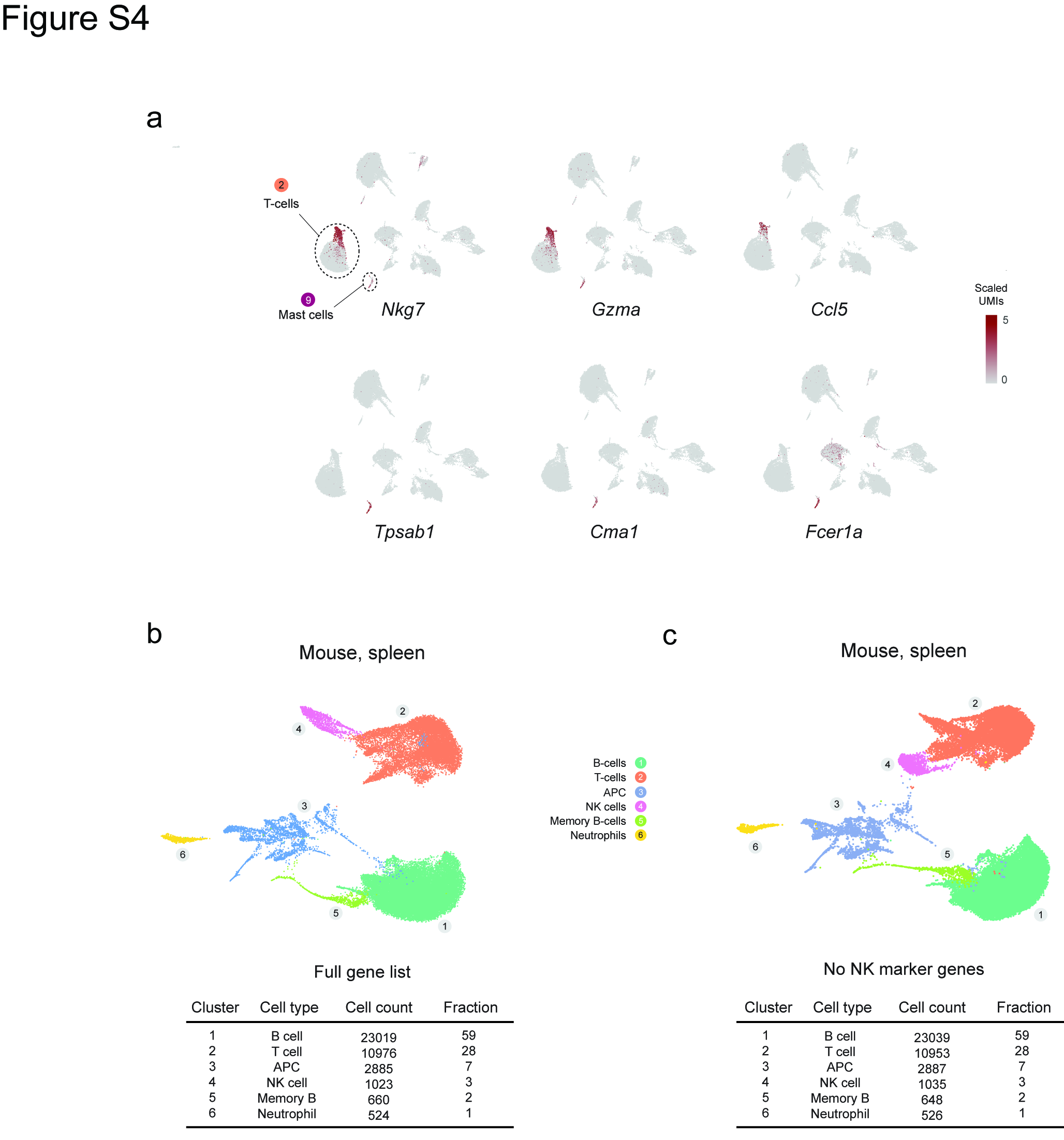

### FigureS5

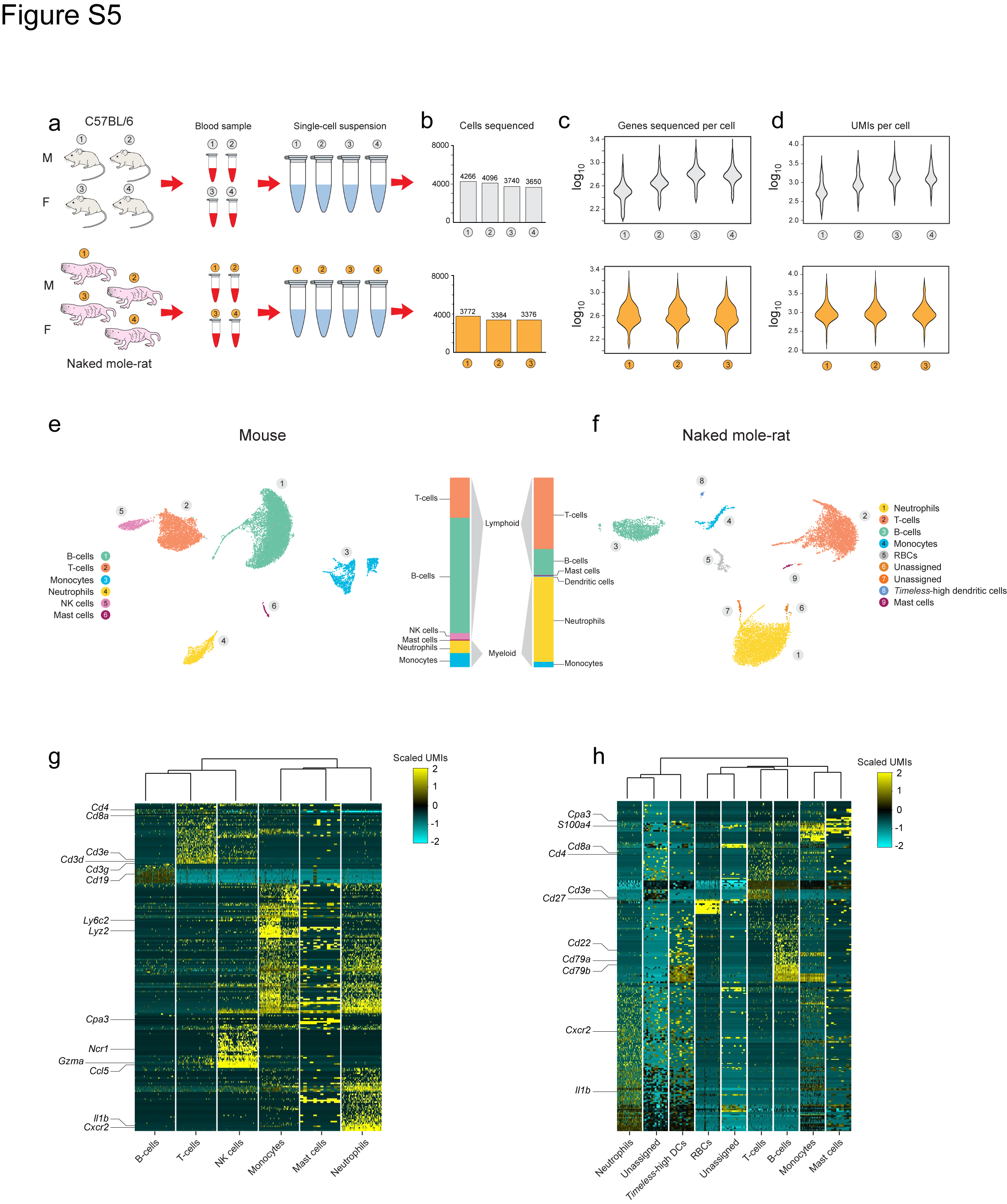

### FigureS6

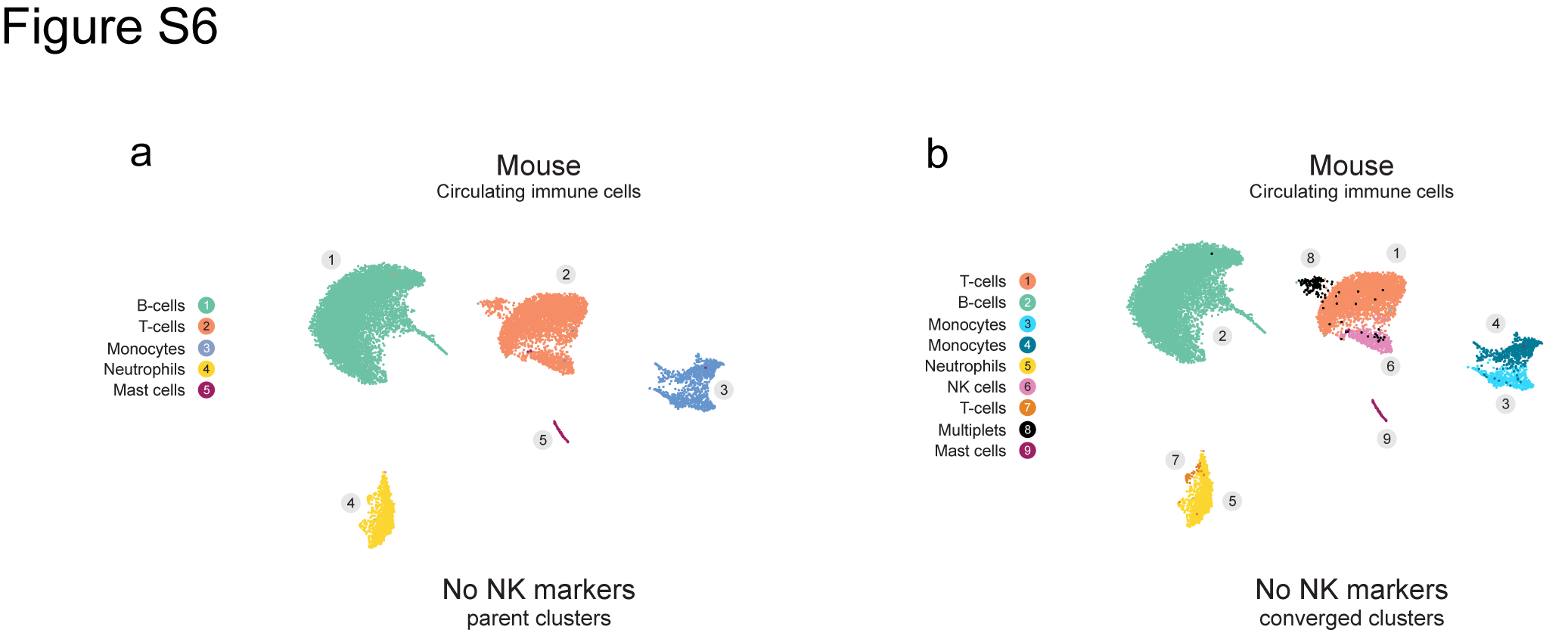

### FigureS8

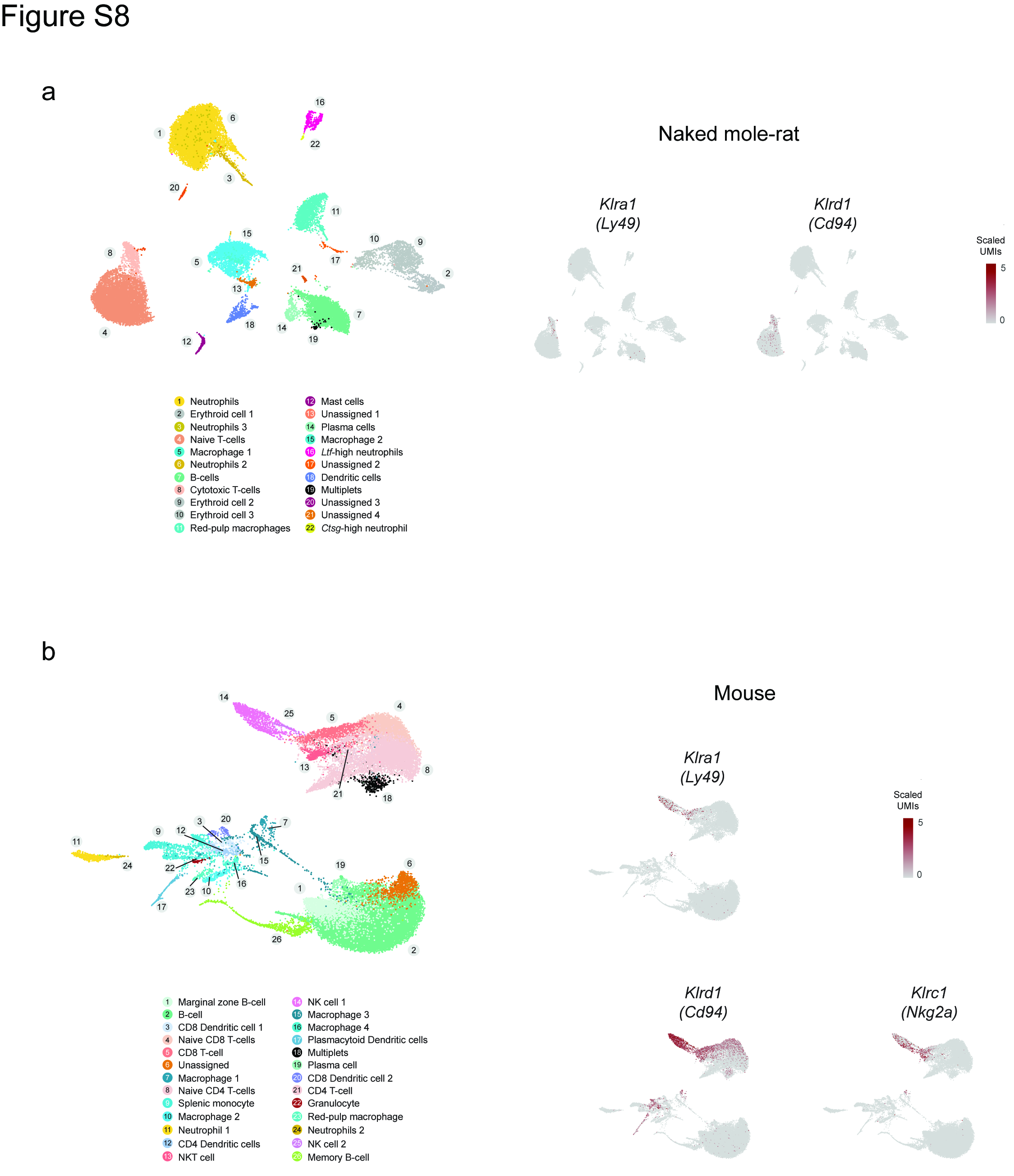

### FigureS9

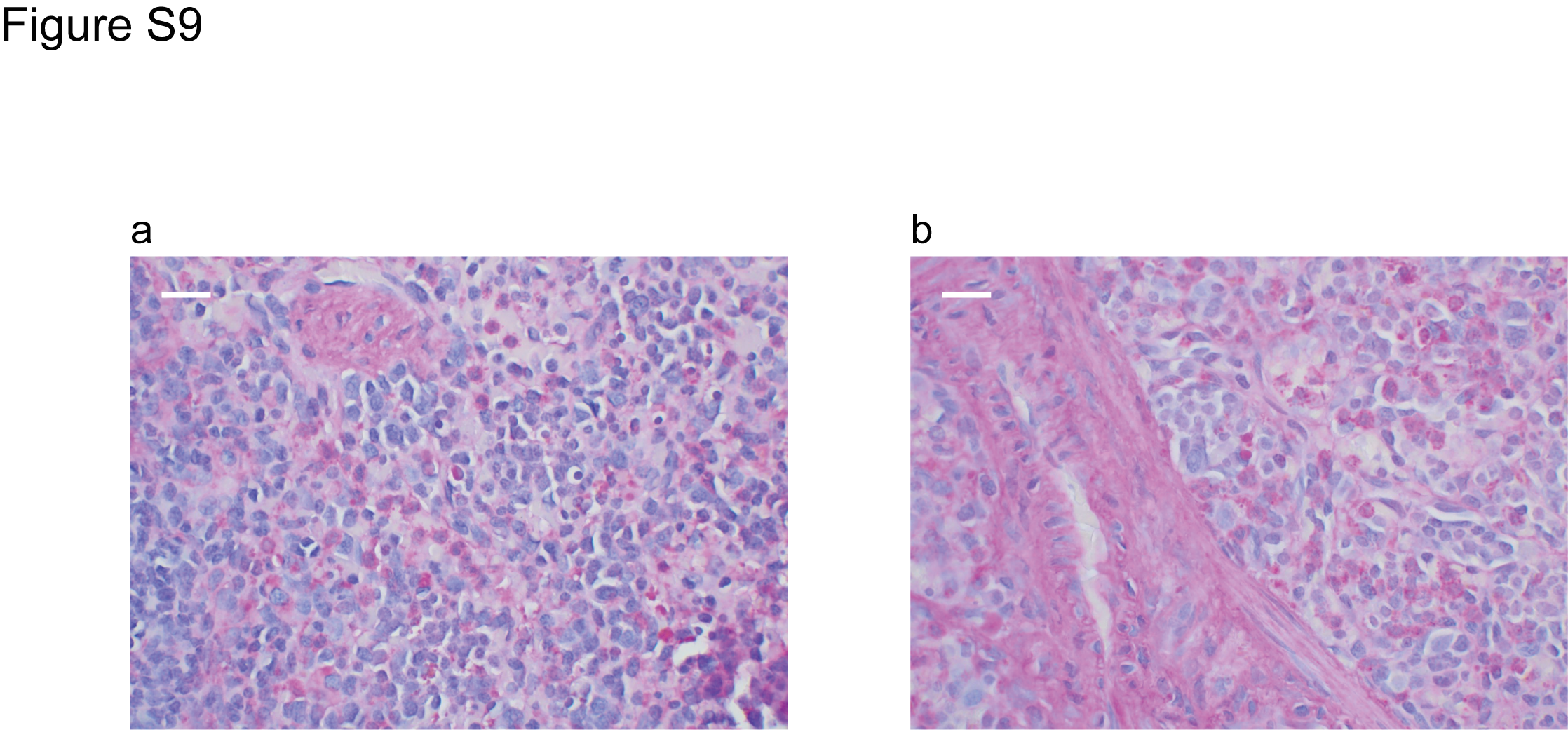

### FigureS10

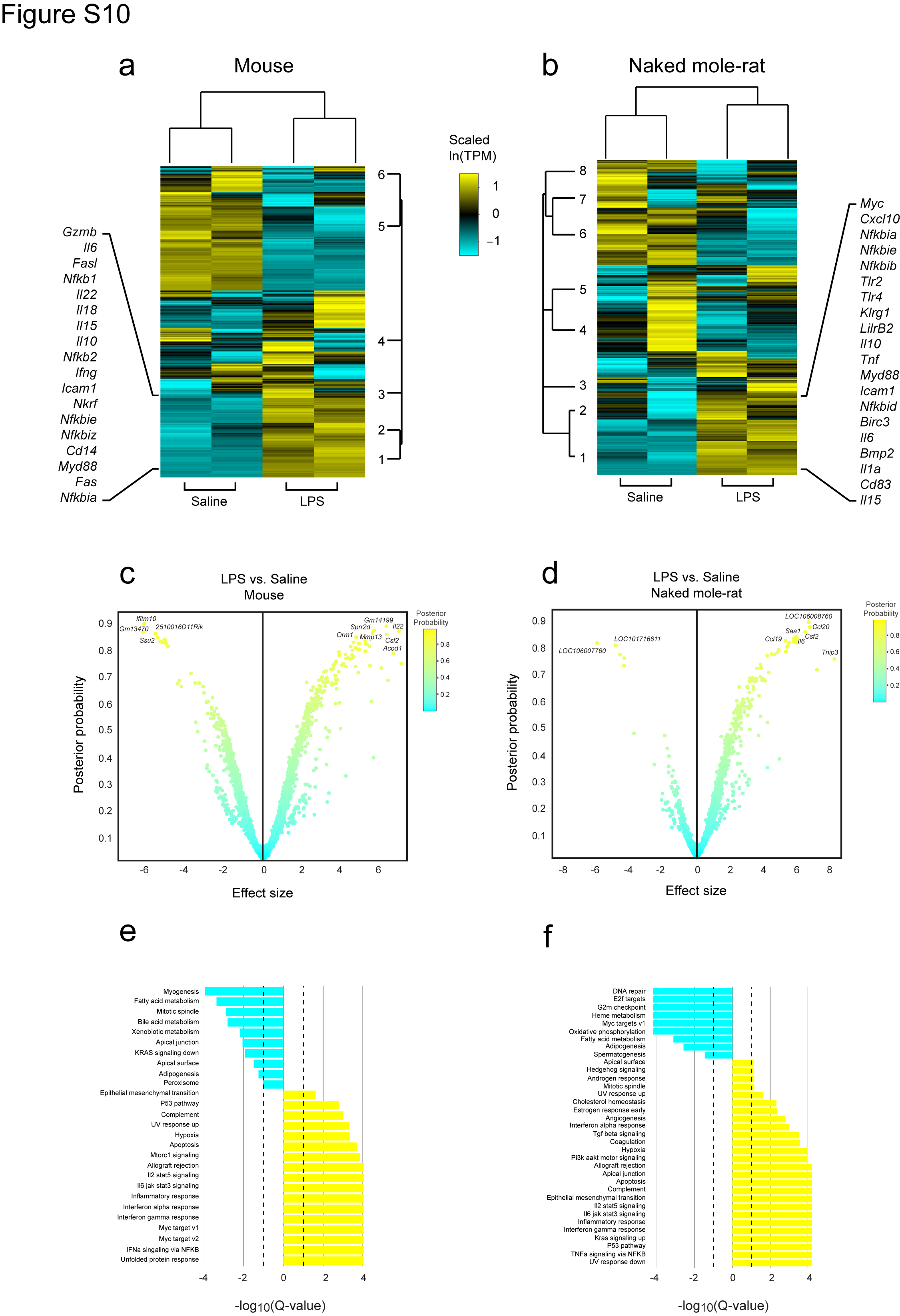

### FigureS11

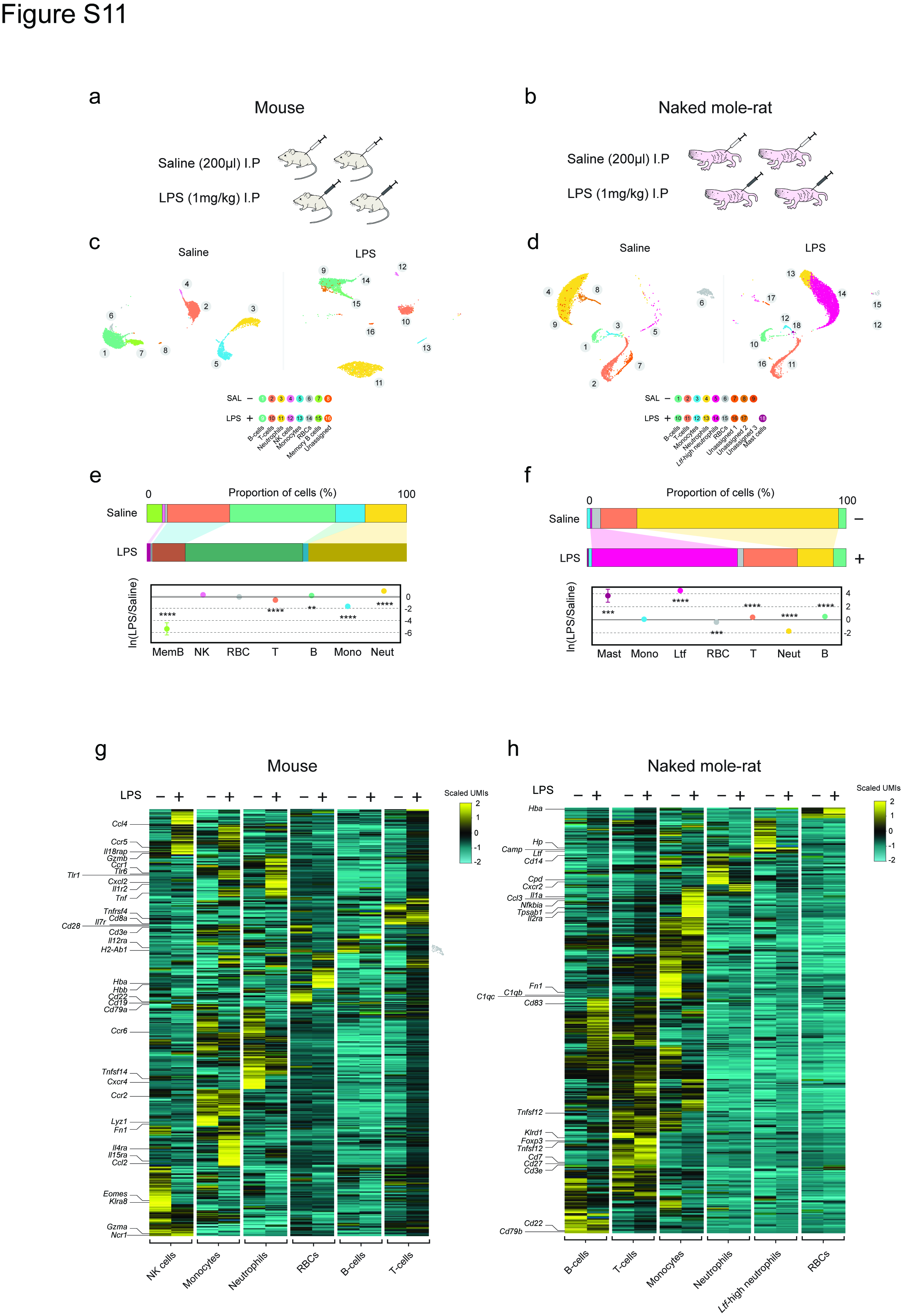

### FigureS12

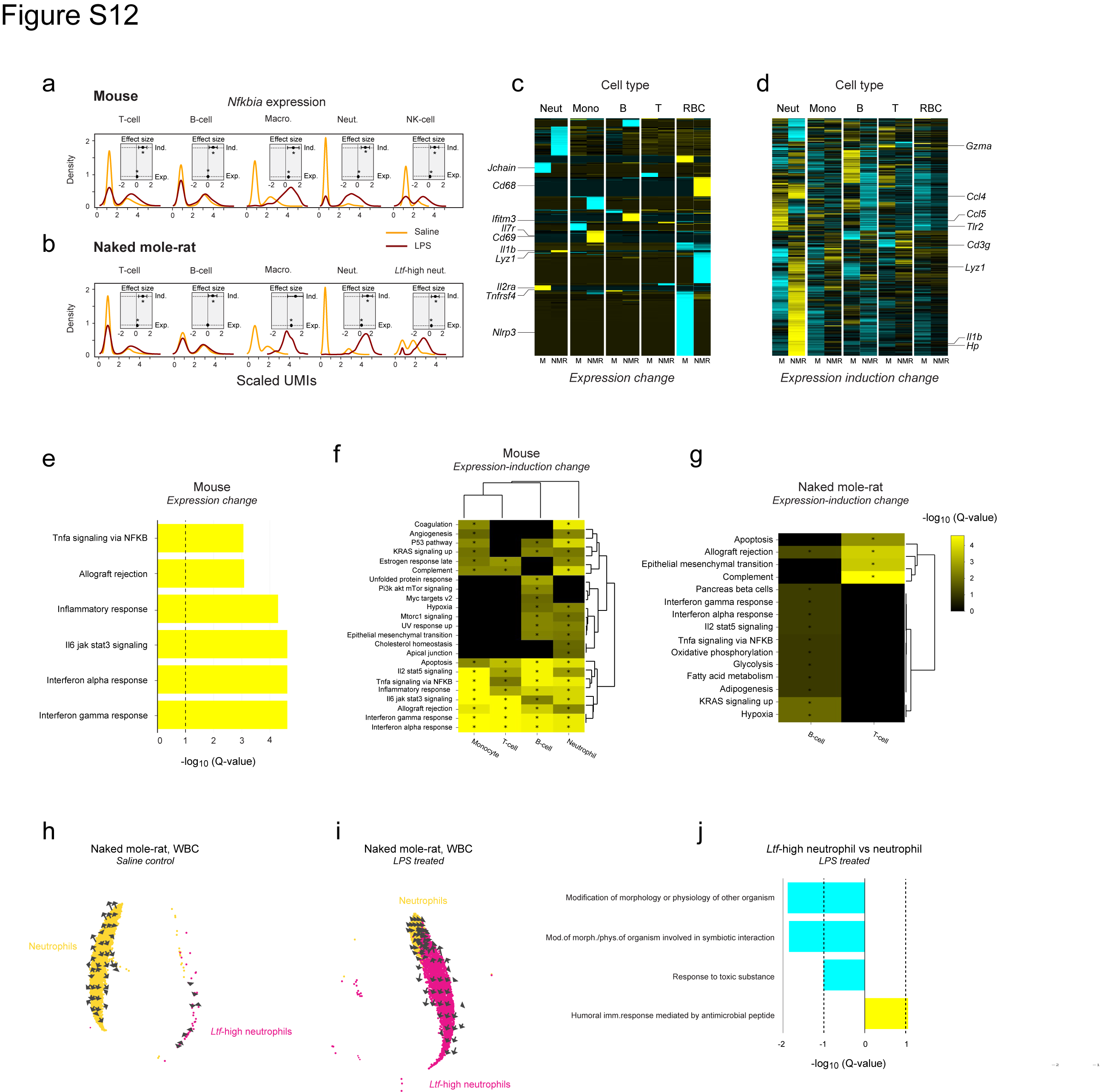
