## Supplementary material for "Single-cell transcriptomics of the naked mole-rat reveals unexpected features of mammalian immunity": FigureS7

### Figure S7

**EXTRACELLULAR REGION**
**INTRACELLULAR REGION**
**TRANSMEMBRANE REGION**
**IMMUNORECEPTOR TYROSINE INHIBITORY MOTIF (ITIM)**
**DOUBLE CYSTEINE RESIDUE**

#### a. *Klra*

>XP_004845991.1 PREDICTED: killer cell lectin-like receptor 2 [Heterocephalus glaber]
MSAEE**V**T**Y**TT**L**RFLQSPSEAQNKLRPNATKGPREDDGKGFAVPWHLIAATLGIICLLLLMTVVVLVMEIFQEKNEKEKIRENLWQA
QNDSSLKERLTNKTLEYDILKNEMHQKNTELELSSKKKCHRKNKTVSKPLQNKGKICEDDLTCCGVKCYYFIKESKDWNGCKLTCQ
DYGLSPVKIDDRDELDFLRLQLGQCYYWIGLSFDTGENRWKWTESGASPGINVRTMLPSSWEGKCAFLSATRISNIACSKTYNCIC
EKRMDGTFPASVCSEEERYPGVAVPGDRIRRVHVAAMDAREAAGITENGFLPCTQG

## b. *Cd94*

>XP_021114355.1 natural killer cells antigen CD94-like, partial [Heterocephalus glaber]
MNEEPRTFPTLNKNSTVKHKRKKDIKNKRSSKELPVITKEPKHHKHKKHTTTADNITSKEDSSHLPWRLI
SSVLGVMFLLLMAGTIVVAVFTANASPETFPTIQQEGPHCQPCPKDWVWFRCSCYYFSMEKLTWSKSQHV
CLSLNASLVKINREEMNFFSLKSFFWTGIYYKKIKNQWYWEN

No double cysteine residue (like bats)

>XP_021114356.1 natural killer cells antigen CD94 [Heterocephalus glaber]
MKKRRKLLPVSD**CC**SCQEKWIGYQCNCYFISDERKTWEESSQLCASHNSSLLQLQNSDELVFMNTNPKFY
WMGLSSAKECAAWLWEDGSALSQNLLPLLPTPSPGYCIAYSPRKVTTSELCGIKNHYMCKQRPI

No transmembrane region

#### c. *Nkg2*

>XP_021114357.1 NKG2-D type II integral membrane protein [Heterocephalus glaber]
MVILLMSPLFLGYIVAVVMAIHFLVMVGIWATVFINFNGKGKLLFSIKSAIVWMCTKLTIYLYIFLESYC
GPCPKNWICYRNSCYQFSNESKNWYQSKASCQSQNSSLLKIYSKVDQDFFKFMKSYHWMGLEQIPENRSW
HWEDVSVLSPDQLSMVEMQNGTCAVYASSFKGYTENCSTPNTYICMQQPI

2 transmembrane domains
No ITIM
No R in transmembrane

>XP_021114358.1 NKG2-D type II integral membrane protein-like [Heterocephalus glaber]
MSDRRAIYSELHPDKVLKKPPRGLSRPETSISINDREKIDPQLNLSPPCFVIRAPLVLADFPSPPEKIFT
ALLGIILLISKASVLAAMVAVLLCKICGGHLTCYGVKCYYFIKESKDWNGCKLTCQDYGLSPVKIDDRDE
LDFLRLQLGQCYYWIGLSSDTGQNGWKWTESSASPGINARTMLPSSWEGKCAFLRETRISYIACSKTYHC
ICEKRMDATFPASVCSEEESLLWKQGELSPIRKEMFLWNLT

No ITIM
No R in transmembrane

>XP_004845990.1 LOW QUALITY PROTEIN: NKG2-A/NKG2-B type II integral membrane protein [Heterocephalus glaber]

MSNHRVVYTELNLAKDPKQQQRKPKGNKSSILESEQQ**I**T**Y**AT**L**TLQSAAQEQRGNGKDHSCKDFTSPPEK
LLAGALGGSCLILVTTVIVVMTVVIPSSVIQEKNKSSLLRTPKAYHCGPCPKKXLTFSDNCYYFGVEKKT
WTESLVYCTNKNSSLLYIDNEEEMKFLGSLSYSSWIGVFRSGIDRPWLWVNGSRVKESSSHEEDCAWLSS
SGLIADNCRSAHTYNCKHKV
